# A large-sample crisis? Exaggerated false positives by popular differential expression methods

**DOI:** 10.1101/2021.08.25.457733

**Authors:** Yumei Li, Xinzhou Ge, Fanglue Peng, Wei Li, Jingyi Jessica Li

## Abstract

We report a surprising phenomenon about identifying differentially expressed genes (DEGs) from population-level RNA-seq data: two popular bioinformatics methods, DESeq2 and edgeR, have unexpectedly high false discovery rates (FDRs). Via permutation analysis on an immunotherapy RNA-seq dataset, we observed that DESeq2 and edgeR identified even more DEGs after samples’ condition labels were randomly permuted. Motivated by this, we evaluated six DEG identification methods (DESeq2, edgeR, limma-voom, NOISeq, dearseq, and the Wilcoxon rank-sum test) on population-level RNA-seq datasets. We found that the FDR control was often failed by the three popular parametric methods—DESeq2, edgeR, and limma-voom— and the new non-parametric method dearseq. In particular, the actual FDRs of DESeq2 and edgeR sometimes exceeded 20% when the target FDR threshold was only 5%. Although NOISeq, a non-parametric method used by GTEx, controlled the FDR better than the other four methods did, its power was much lower than that of the Wilcoxon rank-sum test, a classic nonparametric test that consistently controlled the FDR and achieved good power in our evaluation. Based on these results, for population-level RNA-seq studies, we recommend the Wilcoxon rank-sum test.

## Main text

RNA-seq is an approach to transcriptome profiling using deep-sequencing technologies^1–3^. Since RNA-seq was developed more than one decade ago, it has become an indispensable tool for genome-wide transcriptomic studies. One primary research task in these studies is the identification of DEGs between two conditions (e.g., tumor and normal samples)^2^. A long-standing, core challenge in this task is the small sample size, which is typically two or three replicates per condition. Many statistical methods have been developed to address this issue by making parametric, restrictive distributional assumptions on RNA-seq data, and the two most popular methods of this type are DESeq2^4^ and edgeR^5^. However, as sample sizes have become large in population-level RNA-seq studies, where dozens to thousands of samples were collected from individuals^6,7^, a natural question to ask is whether DESeq2 and edgeR remain appropriate.

To evaluate the performance of DESeq2 and edgeR, we applied both methods to 13 population-level RNA-seq datasets with total sample sizes ranging from 100 to 1376 (**Supplementary Table 1**). We found that DESeq2 and edgeR had large discrepancies in the DEGs they identified on these datasets (**Supplementary Fig. 1**). In particular, 23.71%–75% of the DEGs identified by DESeq2 were missed by edgeR. The most surprising result is from an immunotherapy dataset (including 51 pre-nivolumab and 58 on-nivolumab anti-PD-1 therapy patients)^8^: DESeq2 and edgeR had only an 8% overlap in the DEGs they identified (DESeq2 and edgeR identified 144 and 319 DEGs, respectively, with a union of 427 DEGs but only 36 DEGs in common). This phenomenon raises a critical question: did DESeq2 and edgeR reliably control their false discovery rates (FDRs) to the target 5% on this dataset?

To answer this question, we generated 668 negative-control datasets by randomly permuting the two-condition labels (pre-nivolumab and on-nivolumab) of the 109 RNA-seq samples in this immunotherapy dataset (**Methods**). Since any DEGs identified from these permuted datasets are known as false positives, we used these permuted datasets to evaluate the FDRs of DESeq2 and edgeR. Surprisingly, DESeq2 and edgeR identified more DEGs from 84.88% and 78.89% of these permuted datasets than from the original dataset (**Fig. 1A**). In particular, DESeq2 and edgeR mistakenly identified 16,791 and 13,448 genes as DEGs, respectively, from at least one permuted dataset (**Fig. 1B**). Even more, as many as 56.25% and 89.34% of the DEGs, which DEseq2 and edgeR identified from the original dataset, were also identified as DEGs from at least one permuted dataset, suggesting that these DEGs were spurious (**Fig. 1C**). These results raise the caution about exaggerated false positives by DESeq2 and edgeR on the original dataset.

**Fig. 1.**
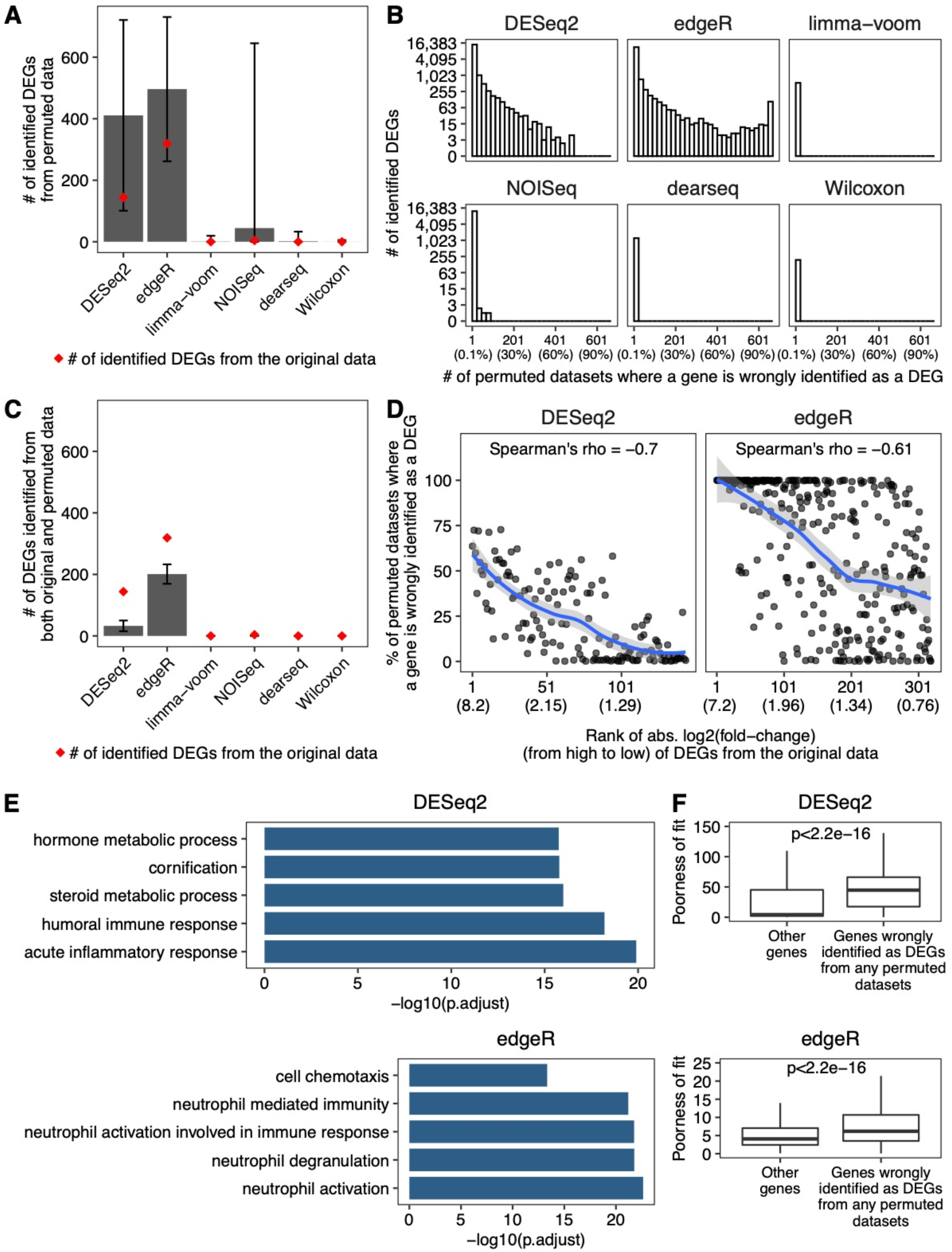
Exaggerated false DEGs identified by DESeq2 and edgeR from anti-PD-1 therapy RNA-seq datasets. **A.** Barplot showing the average numbers of DEGs identified from 668 permuted datasets. The error bars represent the standard deviations of 668 permutations. The red dots indicate the numbers of DEGs identified from the original dataset. **B.** The distributions of the number of permuted datasets where a gene was mistakenly identified as a DEG. The percentages corresponding to the numbers are listed in parentheses below the numbers. **C.** Barplot showing the average numbers of DEGs identified from both the original dataset and the permuted datasets. The error bars represent the standard deviations of 668 permutations. The red dots indicate the numbers of DEGs identified from the original dataset. **D.** Percentage of permuted datasets where a DEG identified from the original dataset was also identified as a DEG. The genes are sorted by absolute log2(fold-change) in the original dataset in decreasing order. The absolute log2(fold-change) values corresponding to the ranks are listed in parentheses below the ranks. The line is fitted using the loess method, and the shaded areas represent 95% confidential intervals. **E.** GO term enrichment for the DEGs identified from at least 10% permuted datasets. The top 5 enriched biological processes GO terms are shown. The analyses were performed using R package clusterProfiler. P.adjust represents the adjusted p-value using the Benjamini & Hochberg method. **F.** Boxplots showing the poorness of fitting the negative binomial model to the genes identified by DESeq2 or edgeR as DEGs from any permuted datasets vs. all the other genes. The poorness of fit for each gene is defined as its negative log10(P-value) from the goodness-of-fit test for the negative binomial distributions estimated by DESeq2 or edgeR. The p-value in each panel was calculated by the Wilcoxon rank-sum test to compare the two groups of genes’ poorness-of-fit values.

What’s more counter-intuitive, the genes with larger fold changes estimated by DESeq2 and edgeR (between the two conditions in the original dataset) were more likely identified as DEGs by the two methods from the permuted datasets (**Fig. 1D and Supplementary Fig. 2**). As biologists tend to believe that these genes are more likely true DEGs (which is not necessarily true because a dataset may contain no true DEGs at all), the fact that these genes are false positives would waste experimental validation efforts.

Out of curiosity and as a means of verification, we investigated the biological functions of the spurious DEGs identified by DESeq2 or edgeR from the permuted datasets. Unexpectedly, these spurious DEGs were enriched in immune-related gene ontology (GO) terms (**Fig. 1E**). Hence, if these spurious DEGs were not removed by FDR control, they would mislead researchers to believe that there was an immune response difference between pre-nivolumab and on-nivolumab patients, a surely undesirable consequence that DEG analysis must avoid.

Then a question follows: why did DESeq2 and edgeR make so many false positive discoveries from this immunotherapy dataset? Our immediate hypothetical reason was the violation of the negative binomial model assumed by both DESeq2 and edgeR^9^. To check this hypothesis, we separated all genes into two groups: (1) the genes identified as DEGs from any permuted datasets and (2) all the other genes; then we evaluated how well the negative binomial model fit to the genes in each group. In line with our hypothesis, the model fitting was worse for the genes in the first group, consistent with the fact these genes were spurious DEGs (**Fig. 1F and Supplementary Fig. 3**).

Motivated by these findings, we further benchmarked DESeq2 and edgeR along with four other representative DEG identification methods on this immunotherapy dataset and the other 12 population-level RNA-seq datasets from the Genotype-Tissue Expression (GTEx) project^7^ and the Cancer Genome Atlas (TCGA)^6^ (**Supplementary Table 1**). The four representative methods include two popular methods limma-voom^10,11^ and NOISeq^12^, a new method dearseq^13^ (which claimed to overcome the FDR control issue of DESeq2 and edgeR on large-sample-size data), and the classic Wilcoxon rank-sum test^14^. Note that DESeq2, edgeR, and limma-voom are parametric methods that assume parametric models for data distribution, while NOISeq, dearseq, and the Wilcoxon rank-sum test are non-parametric methods that are less restrictive but require large sample sizes to have good power. (The GTEx project used DESeq2 and NOISeq for DEG identification.) Using permutation analysis on these datasets, we found that DESeq2 and edgeR consistently showed exaggerated false positives (reflected by their underestimated FDRs) compared to the other four methods (**Supplementary Fig. 4-15**).

While the permutation analysis created true negatives (non-DEGs) to allow FDR evaluation, it did not allow the evaluation of DEG identification power, which requires true positives (DEGs) to be known. Hence, we generated 50 (identically and independently distributed) semi-synthetic datasets with known true DEGs and non-DEGs from each of the 12 GTEx and TCGA datasets. Then we used these semi-synthetic datasets to evaluate the FDRs and power of the six DEG identification methods (**Methods**). In the comparison between 386 heart left ventricle samples and 372 atrial appendage samples in a GTEx dataset, only the Wilcoxon rank-sum test consistently controlled the FDR under a range of thresholds from 0.001% to 5% (**Fig. 2A**). In contrast, the other five methods, especially DESeq2 and edgeR, failed to control the FDR consistently. Moreover, we compared the power of the six methods conditional on their actual FDRs (**Methods**). (Due to the tradeoff between FDR and power, power comparison is only valid when FDR is equal.) As shown in **Fig. 2A**, the Wilcoxon rank-sum test outperformed the other five methods in terms of power.

**Fig. 2.**
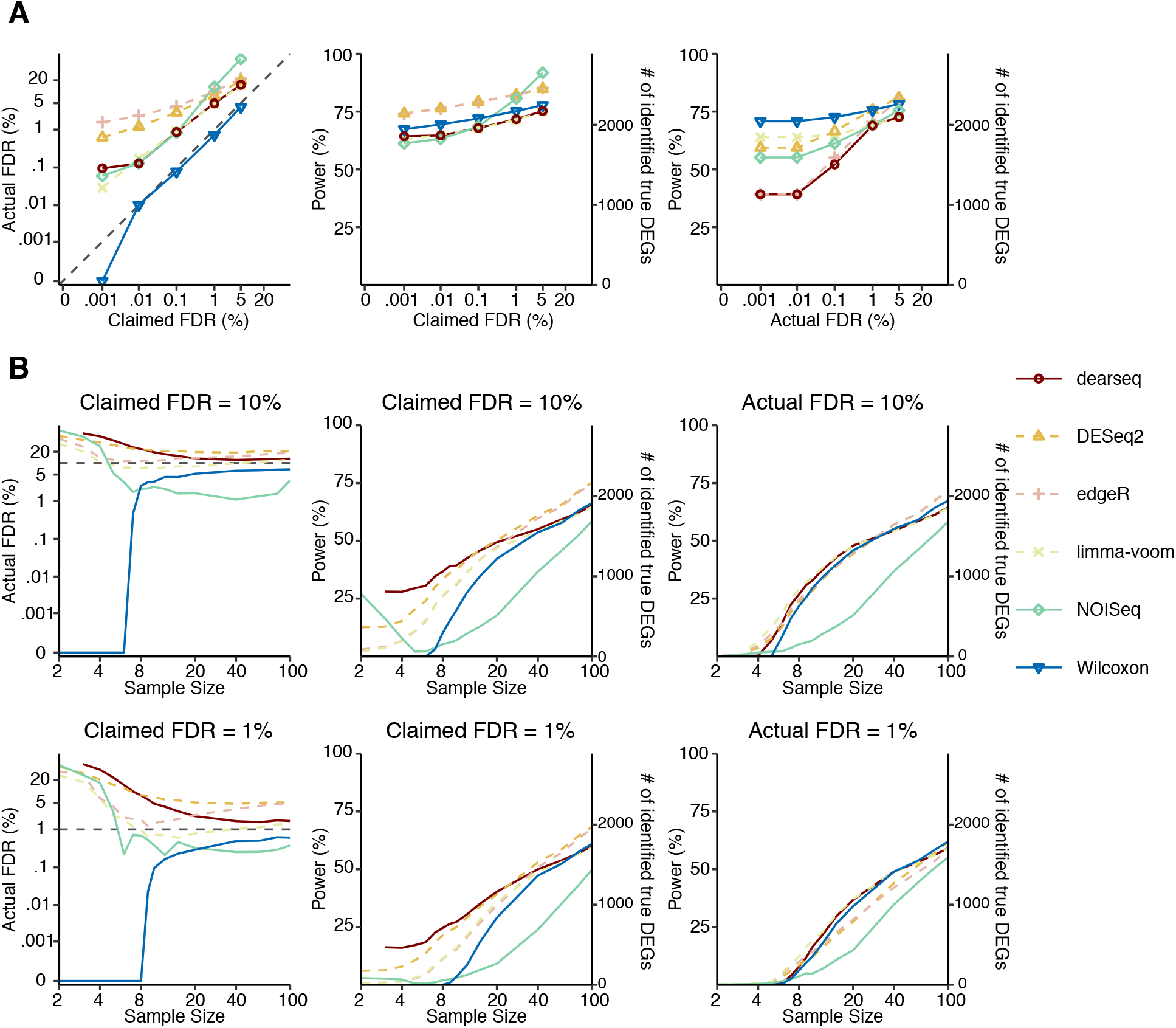
Wilcoxon test has the best FDR control and power on heart left ventricle vs. atrial appendage GTEx datasets with synthetic ground truths. **A.** The FDR control (left panel), power (middle panel) given the claimed FDRs, and power given the actual FDRs (right panel) under a range of FDR thresholds from 0.001% to 5%. **B.** The FDR control (left), power given the claimed FDRs (middle), and power given the actual FDRs (right) for a range of per-condition sample sizes from 2 to 100, under FDR thresholds 10% (top panels) and 1% (bottom panels). The claimed FDRs, actual FDRs, and power were all calculated as the averages of 50 randomly down-sampled datasets.

Finally, to investigate how sample sizes influence the performance of the six methods, we down-sampled each semi-synthetic dataset to obtain per-condition sample sizes ranging from 2 to 100. Again, only the Wilcoxon rank-sum test consistently controlled the FDR at all sample sizes (**Fig. 2B**). Granted, at the FDR threshold 1%, the Wilcoxon rank-sum test had almost no power when the per-condition sample size was smaller than 8—an expected phenomenon for its nonparametric nature. However, when the per-condition sample size exceeded 8, the Wilcoxon rank-sum test achieved comparable or better power compared with the three parametric methods (DESeq2, edgeR, and limma-voom) and the new method dearseq, and it clearly outpowered NOIseq (**Fig. 2B**). These observations were consistent across all 600 semi-synthetic datasets **(Supplementary Figs. 16-26**). In summary, when the per-condition sample size is less than 8, parametric methods may be used because their power advantage may outweigh their possibly exaggerated false positives; however, for large-sample-size data, the Wilcoxon rank-sum test is the best choice for its solid FDR control and good power.

The three parametric methods—DESeq2, edgeR, and limma—have long been dominant in transcriptomic studies. For example, the GTEx project, a consortium effort studying gene expression and regulation in normal human tissues, used DESeq2 coupled with NOISeq to find DEGs between tissues^15^; several studies applied edgeR or limma to TCGA RNA-seq data to find DEGs between tumor and normal samples^16–18^; with increasing attention on the immunotherapy, researchers used DESeq2 to detect DEGs between responders and non-responders of the immunotherapy^8,19^. However, while the three parametric methods were initially designed to address the small-sample-size issue, these population-level studies had much larger sample sizes (at least dozens) and thus no longer needed restrictive parametric assumptions. Moreover, violation of parametric assumptions may lead to ill-behaved P-values and thus failed FDR control^20^, an issue independent of the sample size.

In this study, we showed the superiority of the Wilcoxon rank-sum test, a powerful and robust non-parametric test also known as the Mann-Whitney test developed in the 1940s^14,21–24^, for analyzing large-sample-size RNA-seq datasets. The Wilcoxon rank-sum test is known to be especially powerful for skewed distributions^25^, as is the case with gene expression counts measured by RNA-seq. Our results also echo the importance of verifying FDR control by permutation analysis. Beyond RNA-seq data analysis, our study suggests that, for population-level studies with large sample sizes, classic non-parametric statistical methods should be considered as the baseline methods for data analysis and new method benchmarking.

Finally, we note that, unlike DESeq2, edgeR, limma-voom, and dearseq, the Wilcoxon rank-sum test is a non-regression-based method, making it unable to adjust for confounders. Hence, to use the Wilcoxon rank-sum test for DEG identification, researchers must first normalize RNA-seq samples to remove possible confounder effects. Another limitation of the Wilcoxon rank-sum test is that it only applies to two-condition comparisons. To compare more than two conditions on population-level data, we recommend the Kruskal–Wallis test, also known as the one-way ANOVA on ranks^26^.

## Acknowledgements

We thank Jason Sheng Li of Wei Li lab for suggestions on the title. We also thank other members of Wei Li lab and Jingyi Jessica Li lab for helpful discussions.

## Author contributions

Y.L., W.L. and J.J.L. conceived and supervised this project. Y.L. and X.Z. performed the data analysis. Y.L., X.Z., F.P., J.J.L and W.L. interpreted the data and wrote the manuscript.

## Data availability

All the permuted and semi-synthetic datasets used to generated results can be found at Zenodo via https://doi.org/10.5281/zenodo.5241320.

## Code availability

All the codes used to generate results can be found at GitHub via URL https://github.com/xihuimeijing/DEGs_Analysis_FDR.

A tutorial for identifying DEGs using the Wilcoxon rank-sum test can be found at https://rpubs.com/LiYumei/806213.

## Supplementary Materials

Materials and Methods

Supplementary Figs. 1 to 26

Supplementary Table 1

