## Supplementary for "A large-sample crisis? Exaggerated false positives by popular differential expression methods"

**This PDF file includes:**

Materials and Methods
Supplementary Figs. 1 to 26
Supplementary Table 1

### Materials and Methods

#### Tool selection

We selected the three parametric methods for identifying differentially expressed gene (DEGs) from RNA-seq data based on popularity: DESeq2, edgeR, and limma-voom (v 3.44.3) (with 26,924, 20,753, and 12,544 citations, respectively, in Google Scholar as of 16 June 2021). We chose the non-parametric method NOISeq because the Genotype-Tissue Expression (GTEx) consortium used it to identify DEGs between tissues, and we used GTEx RNA-seq datasets in our study. We also chose the newly developed non-parametric method dearseq, which claimed that it overcomes the FDR control issue of DESeq2, edgeR, and limma-voom on large-sample-size data. Moreover, we included the Wilcoxon rank-sum test, a classical non-parametric statistical test about whether two samples (i.e., a gene's two sets of expression levels measured under two conditions) follow the same distribution.

#### Datasets

The RNA-seq datasets used in this study all have at least 50 samples per condition.

- For the immunotherapy study, we selected one dataset with a total sample size of 109, including 51 pre-nivolumab and 58 on-nivolumab anti-PD-1 therapy melanoma samples (https://www.ncbi.nlm.nih.gov/geo/query/acc.cgi?acc=GSE91061).
- For TCGA data, we selected RNA-seq datasets of six cancer types, which have paired normal tissues and sample sizes greater than 50 for both cancer and normal tissues. Then we downloaded the gene read count matrices of these selected datasets from GDC Xena Hub (https://xenabrowser.net/datapages/?hub=https://gdc.xenahubs.net:443, release v18.0).

- For GTEx data, we selected six pairs of tissues with sample sizes ranging from 126 to 706. Then we downloaded the gene read count matrices of these tissue samples from the GTEx Portal (https://storage.googleapis.com/gtex_analysis_v8/rna_seq_data/GTEx_Analysis_2017-06-05_v8_RNASeQCv1.1.9_gene_reads.gct.gz, GTEx Analysis V8).

**Supplementary Table 1** lists the detailed information of the datasets used in this study.

### The identification of DEGs

All six methods (DESeq2, edgeR, limma-voom, NOISeq, dearseq, and the Wilcoxon rank-sum test) took a read count matrix and a condition label vector as input. The parameters were set based on the user guides of these methods' software packages.

- For DESeq2 (v1.28.1), we used the *DESeq* function to perform differential analysis, followed by generating the results using the *results* function.
- For edgeR (v3.30.3), we first filtered out genes with very low counts using the *filterByExpr* function, followed by normalization using the trimmed mean of M values (TMM) method. Then the quasi-likelihood F-test was used for differential analysis.
- For limma-voom (v 3.44.3), we filtered genes and calculated the normalization factor in the same way as we did for edgeR. Then we applied the voom transformation to the normalized and filtered count matrix and performed the differential analysis using the *lmFit* and *eBayes* functions.
- For NOISeq (v2.31.0), we used the *noiseqbio* function to identify DEGs.
- For dearseq, the filtering and normalization steps were the same as those for edgeR. Then we used the *dear_seq* function with the asymptotic test to identify DEGs.

- For the Wilcoxon rank-sum test, the filtering and normalization steps were the same as those for edgeR. For p-value calculation, we input each gene's counts-per-million (CPM) values into the *wilcox.test* function in R (v4.0.2). Then we set a p-value cutoff based on an FDR threshold using the Benjamini & Hochberg method.

Specifically, DEGs were selected based on the corresponding FDR threshold for all the six methods (FDR<0.05 for the immunotherapy dataset; FDR<0.01 for GTEx and TCGA datasets).

**<u>The generation of permuted and semi-synthetic data from the original RNA-seq data</u>**

From each original RNA-seq dataset, we generated permuted datasets between two conditions (pre-therapy and on-therapy samples for the immunotherapy data; two tissue types for GTEx data; normal and tumor samples for TCGA data). We used $\boldsymbol{M}$ to denote a gene-by-sample read count matrix (with genes as rows and samples as columns) and $\boldsymbol{C}$ to denote the vector of sample conditions labels (corresponding to the columns of $\boldsymbol{M}$). Then we generated a permuted dataset by randomly permuting all values in $\boldsymbol{C}$ and keeping the original order of samples in $\boldsymbol{M}$. We repeated this permutation procedure for 1000 times to generate 1000 permuted datasets.

Note that some permuted datasets did not mix the two condition labels well (e.g., only two samples were swapped between the two conditions), so they were too similar to the original dataset and might not satisfy the null hypothesis that each gene's expression levels follow the same distribution. To address this issue, we only preserve the permuted datasets whose fractions of samples having original condition labels were between the $1^{st}$ and $3^{rd}$ quantiles (including the two quantiles calculated by the R function quantile() with default arguments) of all 1000 permuted datasets. With this filtering step, we assumed that all preserved permuted datasets satisfy the null hypothesis, so all genes identified as DEGs from any of them were false positives.

The semi-synthetic datasets were generated based on original RNA-seq samples from GTEx and TCGA. We first used all five DEG identification methods to identify DEGs from each original dataset containing two conditions. We then defined *true DEGs* as the genes identified as DEGs by all five methods at a very small FDR threshold (0.0001%). We used $\boldsymbol{X}$ and $\boldsymbol{Y}$ to denote the read count matrices from the two conditions, and $\boldsymbol{X_i}$ and $\boldsymbol{Y_i}$ to denote the read counts of gene $i$ from the two conditions (i.e., the $i$-th row of $\boldsymbol{X}$ and $\boldsymbol{Y}$). Then we generated semi-synthetic datasets $\boldsymbol{X'}$ and $\boldsymbol{Y'}$ in the following way: we preserved the read counts of true DEGs; for each of the remaining genes, we randomly permuted its read counts between the two conditions. That is, $\boldsymbol{X'_i} = \boldsymbol{X_i}$ and $\boldsymbol{Y'_i} = \boldsymbol{Y_i}$ if gene $i$ is a true DEG, $(\boldsymbol{X'_i}, \boldsymbol{Y'_i}) = \sigma_i(\boldsymbol{X_i}, \boldsymbol{Y_i})$ if gene $i$ is not a true DEG, where $\sigma_i$ is a random permutation of values in $\boldsymbol{X_i}$ and $\boldsymbol{Y_i}$. We repeated this procedure independently for 50 times to generate 50 semi-synthetic datasets. To generate down-sampled semi-synthetic datasets with a per-condition sample size of $n$, we randomly sampled $n$ columns from $\boldsymbol{X'}$ and $\boldsymbol{Y'}$ each.

**Calculation of FDR, power, and poorness of model fit**

The FDR is defined as the expectation of the false discovery proportion (FDP), the proportion of false positives among all the discoveries. The FDR cannot be directly observed, but the FDP can be calculated from benchmark datasets with known true positives and negatives. In our semi-synthetic data analysis, we defined true DEGs as true positives and the remaining genes as true negatives. First, we calculated the FDP of each DEG identification method (e.g., DESeq2) on each semi-synthetic dataset. Second, we calculated the method's (approximate) FDR by taking the average of its FDPs on the 50 semi-synthetic datasets.

The power of a DEG identification method is defined as the probability of identifying a gene as a DEG conditional on that the gene is a true DEG. It can also be considered as the expectation of the empirical power, which is the proportion of true DEGs being identified as DEGs. In our synthetic datasets with true DEGs and true non-DEGs, we calculated the power of a method by taking the average of its empirical power on the 50 synthetic datasets, similar to how we calculated the FDR.

We used the goodness-of-fit test to evaluate how well a gene's read counts under a condition can be fit by the negative binomial models estimated by DESeq2 and edgeR. To remove batch effects, we used the normalized read counts output by DESeq2 and edgeR. For each gene under each condition and each method (DESeq2 or edgeR), we conducted the goodness-of-fit test on the method's normalized read counts, with the method's estimated dispersion parameter of the negative binomial distribution. The goodness-of-fit test was implemented using the function *goodfit* in the R package *vcd* as

```
summary(goodfit(round(normalized_counts), type = "nbinomial", par =
list(size = 1/dispersion)))
```

which returns a p-value. A smaller p-value indicates a poorer fit. Hence, we defined the poorness of fit as the negative $\log_{10}$(p-value).

**Supplementary Fig. 1**

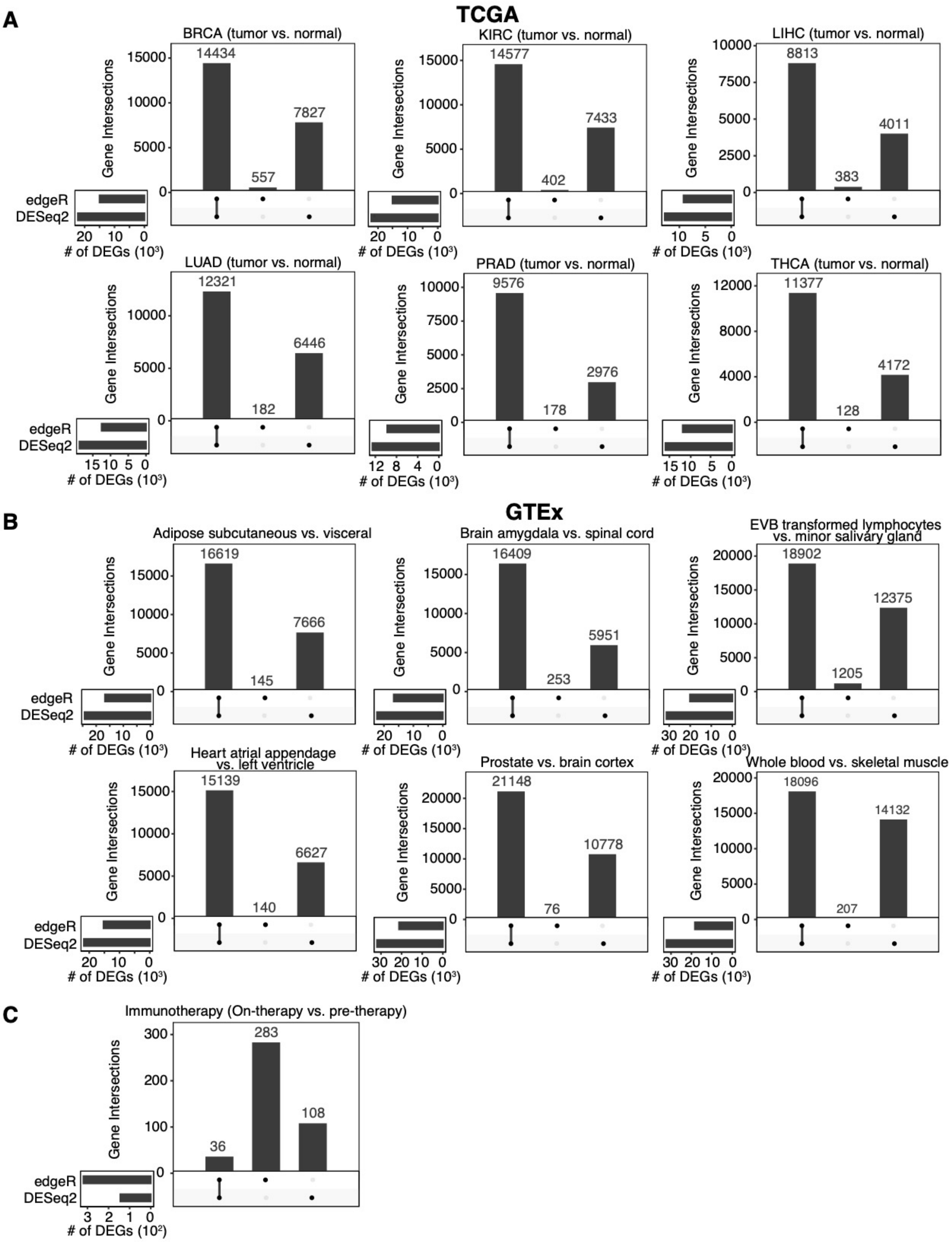

**Supplementary Fig. 1. The large discrepancies in the DEGs identified by DESeq2 and edgeR.**

**(A-C)** Upset plots showing the intersections of the DEGs identified by DESeq2 and edgeR from TCGA (**A**), GTEx (**B**), and immunotherapy (**C**) RNA-seq datasets.

**Supplementary Fig. 2**

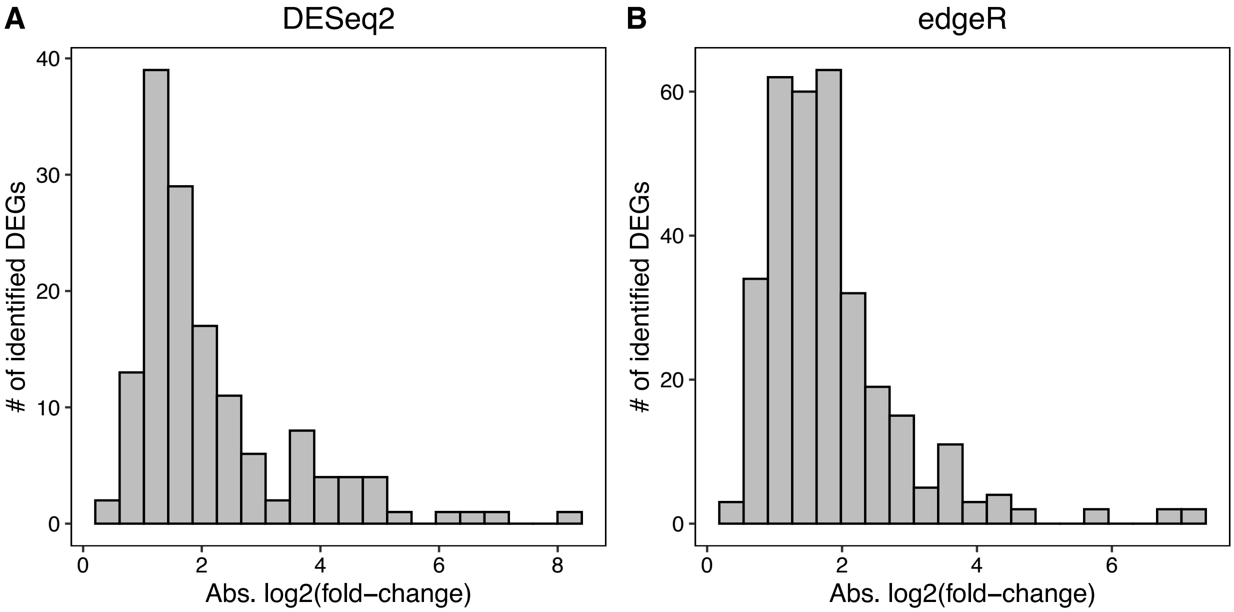


**Supplementary Fig. 2. The distribution of gene expression fold changes between pre-therapy and on-therapy samples in the original immunotherapy dataset calculated by DESeq2 (A) and edgeR (B).**

**Supplementary Fig. 3**

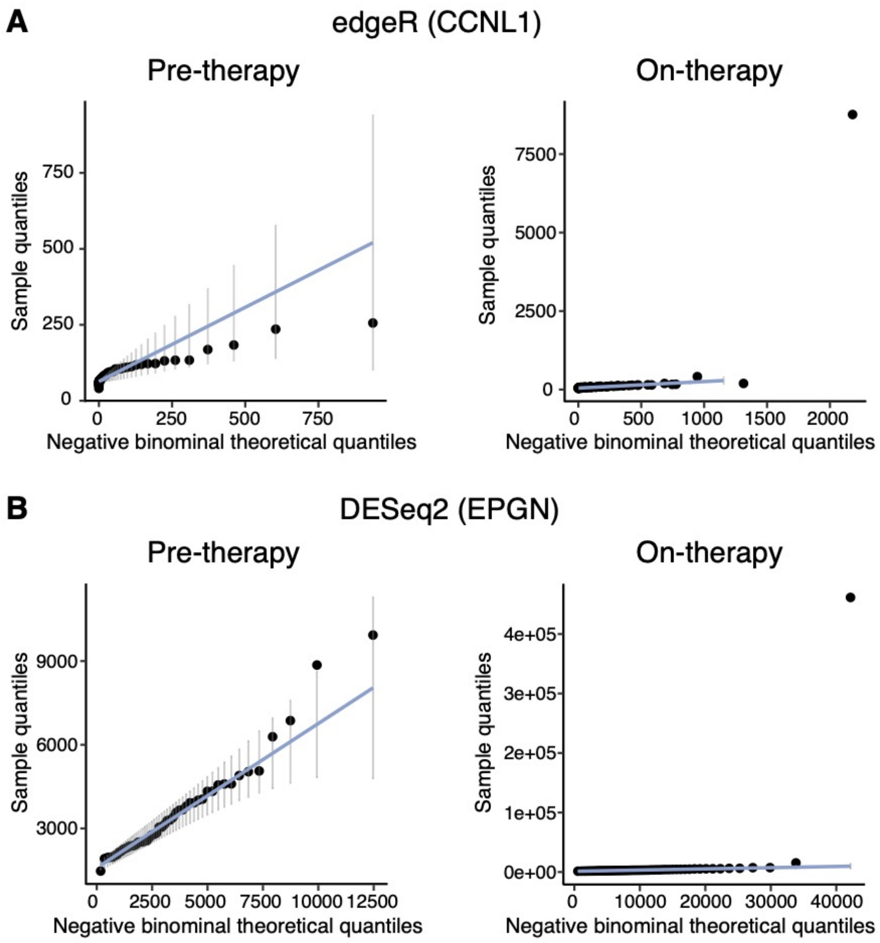


**Supplementary Fig. 3. Quantile-quantile (Q-Q) plots showing the discrepancy between observed read counts and the negative binominal theoretical quantiles estimated by edgeR and DESeq2.**

**A.** The Q-Q plot for *CCNL1* with theoretical read counts in two conditions (pre-therapy and on-therapy) estimated by edgeR.

**B.** The Q-Q plot for *EPGN* with theoretical read counts in two conditions (pre-therapy and on-therapy) estimated by DESeq2.

**Supplementary Fig. 4**

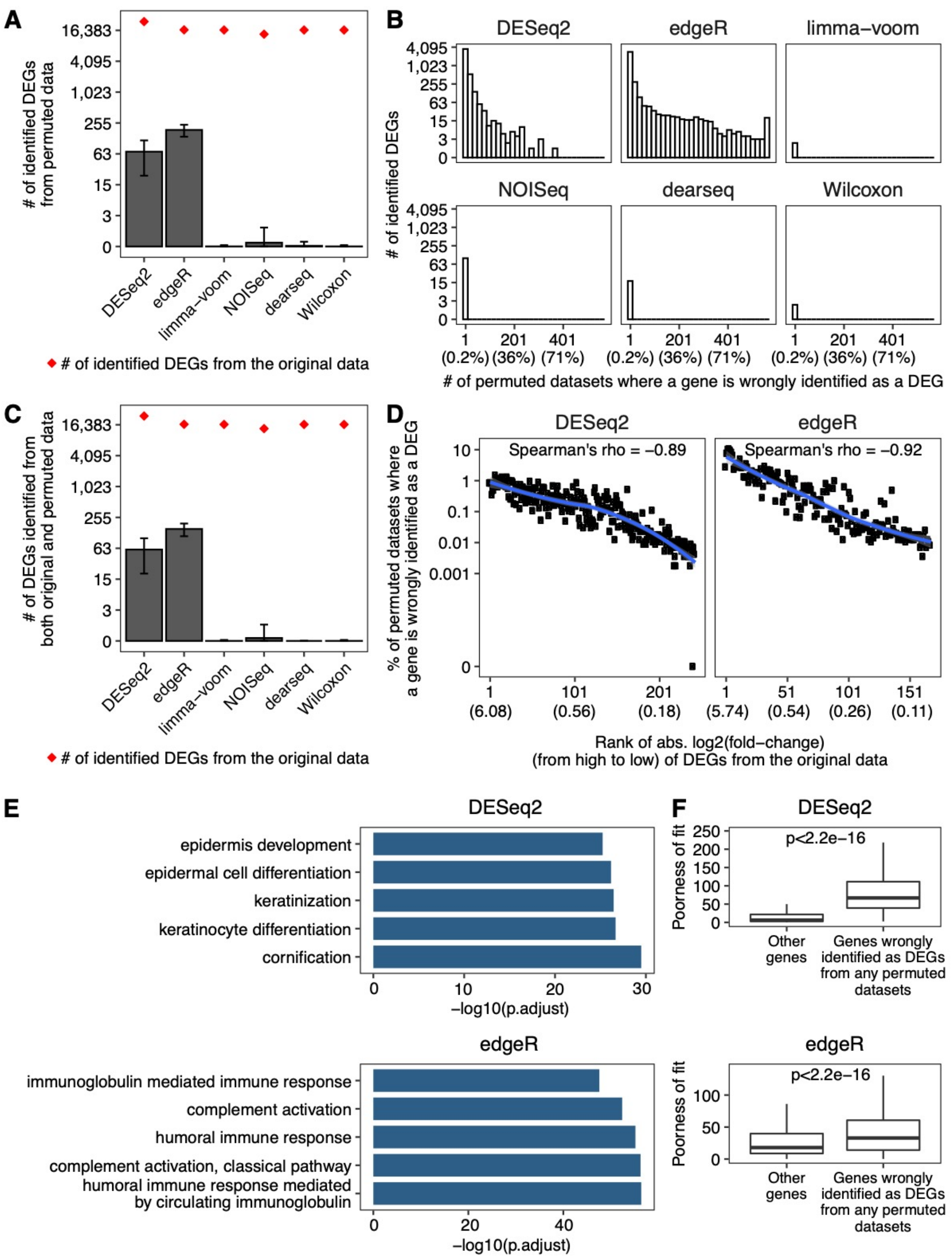


**Supplementary Fig. 4. Exaggerated false DEGs identified by DESeq2 and edgeR from adipose (subcutaneous vs. visceral) GTEx RNA-seq datasets.**

**A.** Barplot showing the average numbers of DEGs identified from 562 permuted datasets. The error bars represent the standard deviations of 562 permutations. The red dots indicate the numbers of DEGs identified from the original dataset.

**B.** The distributions of the number of permuted datasets where a gene was mistakenly identified as a DEG. The percentages corresponding to the numbers are listed in parentheses below the numbers.

**C.** Barplot showing the average numbers of DEGs identified from both the original dataset and the permuted datasets. The error bars represent the standard deviations of 562 permutations. The red dots indicate the numbers of DEGs identified from the original dataset.

**D.** Percentage of permuted datasets where a DEG identified from the original dataset was also identified as a DEG. The genes are sorted by absolute log2(fold-change) in the original dataset in decreasing order and the average values of each 100 genes are shown. The absolute log2(fold-change) values corresponding to the ranks are listed in parentheses below the ranks.The line is fitted using the loess method, and the shaded areas represent 95% confidential intervals.

**E.** GO term enrichment for the DEGs identified from at least 10% permuted datasets. The top 5 enriched biological processes GO terms are shown. The analyses were performed using R package clusterProfiler. P.adjust represents the adjusted p-value using the Benjamini & Hochberg method.

**F.** Boxplots showing the poorness of fitting the negative binomial model to the genes identified by DESeq2 or edgeR as DEGs from any permuted datasets vs. all the other genes. The poorness of fit for each gene is defined as its negative $\log_{10}$(P-value) from the Pearson's chi-squared test for the negative binomial distribution. The p-value in each panel was calculated by the Wilcoxon rank-sum test to compare the two groups of genes' poorness-of-fit values.

**Supplementary Fig. 5**

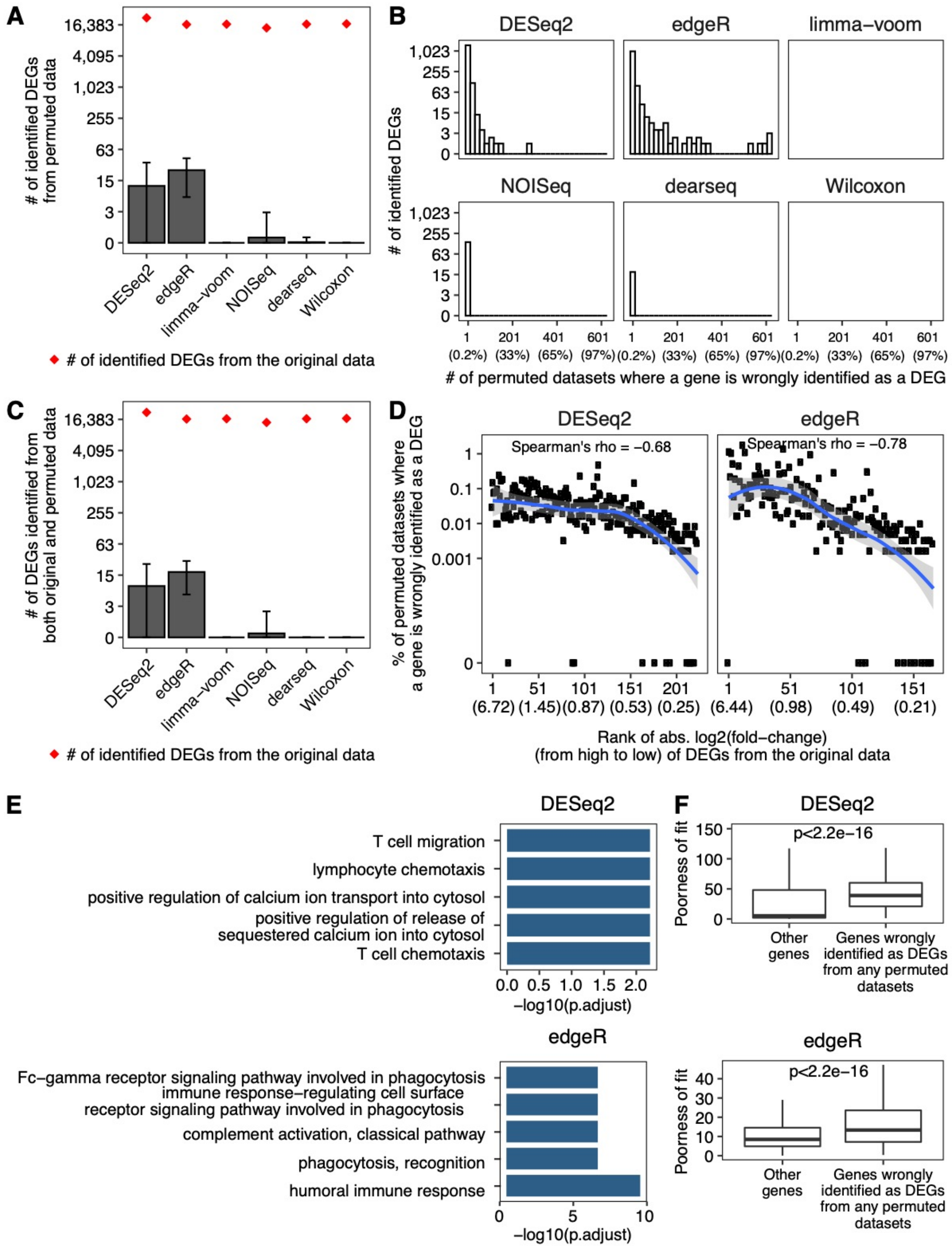


**Supplementary Fig. 5. Exaggerated false DEGs identified by DESeq2 and edgeR from brain (amygdala vs. spinal cord) GTEx RNA-seq datasets.**

**A.** Barplot showing the average numbers of DEGs identified from 617 permuted datasets. The error bars represent the standard deviations of 617 permutations. The red dots indicate the numbers of DEGs identified from the original dataset.

**B.** The distributions of the number of permuted datasets where a gene was mistakenly identified as a DEG. The percentages corresponding to the numbers are listed in parentheses below the numbers.

**C.** Barplot showing the average numbers of DEGs identified from both the original dataset and the permuted datasets. The error bars represent the standard deviations of 617 permutations. The red dots indicate the numbers of DEGs identified from the original dataset.

**D.** Percentage of permuted datasets where a DEG identified from the original dataset was also identified as a DEG. The genes are sorted by absolute log2(fold-change) in the original dataset in decreasing order and the average values of each 100 genes are shown. The absolute log2(fold-change) values corresponding to the ranks are listed in parentheses below the ranks.The line is fitted using the loess method, and the shaded areas represent 95% confidential intervals.

**E.** GO term enrichment for the DEGs identified from at least 10% permuted datasets. The top 5 enriched biological processes GO terms are shown. The analyses were performed using R package clusterProfiler. P.adjust represents the adjusted p-value using the Benjamini & Hochberg method.

**F.** Boxplots showing the poorness of fitting the negative binomial model to the genes identified by DESeq2 or edgeR as DEGs from any permuted datasets vs. all the other genes. The poorness of fit for each gene is defined as its negative $\log_{10}$(P-value) from the Pearson's chi-squared test for the negative binomial distribution. The p-value in each panel was calculated by the Wilcoxon rank-sum test to compare the two groups of genes' poorness-of-fit values.

**Supplementary Fig. 6**

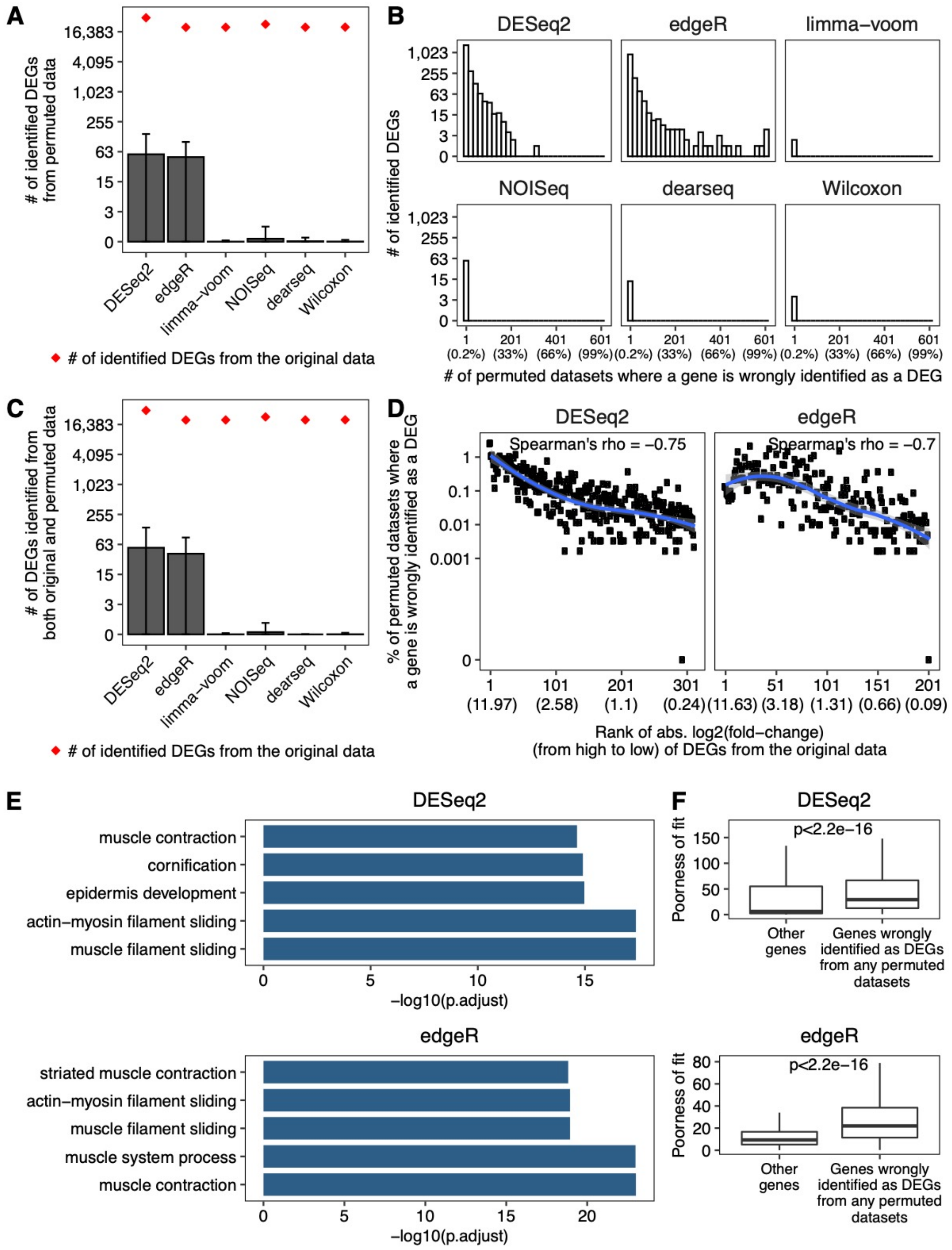


**Supplementary Fig. 6. Exaggerated false DEGs identified by DESeq2 and edgeR from EVB transformed lymphocytes vs. minor salivary gland GTEx RNA-seq datasets.**

**A.** Barplot showing the average numbers of DEGs identified from 607 permuted datasets. The error bars represent the standard deviations of 607 permutations. The red dots indicate the numbers of DEGs identified from the original dataset.

**B.** The distributions of the number of permuted datasets where a gene was mistakenly identified as a DEG. The percentages corresponding to the numbers are listed in parentheses below the numbers.

**C.** Barplot showing the average numbers of DEGs identified from both the original dataset and the permuted datasets. The error bars represent the standard deviations of 607 permutations. The red dots indicate the numbers of DEGs identified from the original dataset.

**D.** Percentage of permuted datasets where a DEG identified from the original dataset was also identified as a DEG. The genes are sorted by absolute log2(fold-change) in the original dataset in decreasing order and the average values of each 100 genes are shown. The absolute log2(fold-change) values corresponding to the ranks are listed in parentheses below the ranks.The line is fitted using the loess method, and the shaded areas represent 95% confidential intervals.

**E.** GO term enrichment for the DEGs identified from at least 10% permuted datasets. The top 5 enriched biological processes GO terms are shown. The analyses were performed using R package clusterProfiler. P.adjust represents the adjusted p-value using the Benjamini & Hochberg method.

**F.** Boxplots showing the poorness of fitting the negative binomial model to the genes identified by DESeq2 or edgeR as DEGs from any permuted datasets vs. all the other genes. The poorness of fit for each gene is defined as its negative $\log_{10}$(P-value) from the Pearson's chi-squared test for the negative binomial distribution. The p-value in each panel was calculated by the Wilcoxon rank-sum test to compare the two groups of genes' poorness-of-fit values.

**Supplementary Fig. 7**

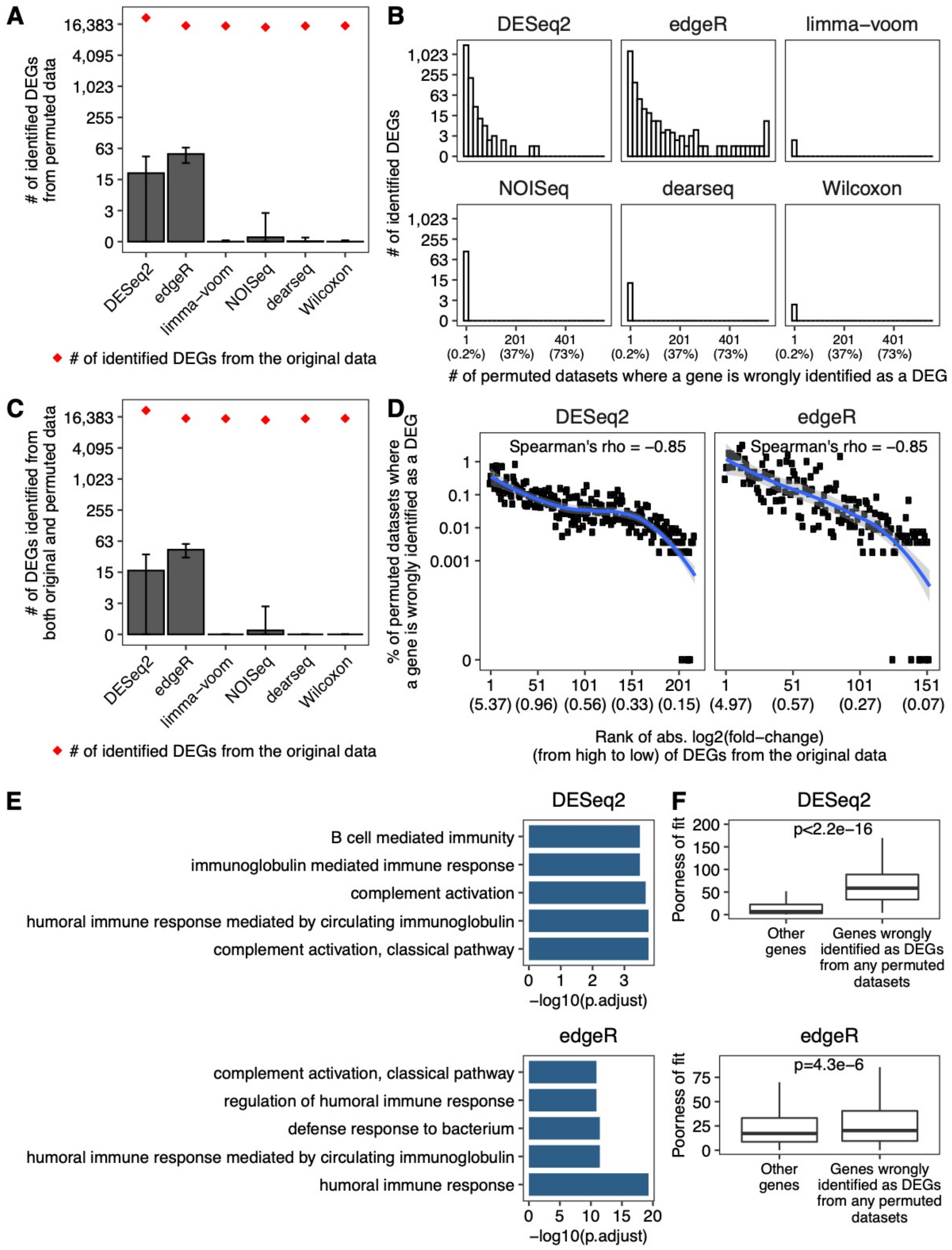


**Supplementary Fig. 7. Exaggerated false DEGs identified by DESeq2 and edgeR from heart (atrial appendage vs. left ventricle) GTEx RNA-seq datasets.**

**A.** Barplot showing the average numbers of DEGs identified from 550 permuted datasets. The error bars represent the standard deviations of 550 permutations. The red dots indicate the numbers of DEGs identified from the original dataset.

**B.** The distributions of the number of permuted datasets where a gene was mistakenly identified as a DEG. The percentages corresponding to the numbers are listed in parentheses below the numbers.

**C.** Barplot showing the average numbers of DEGs identified from both the original dataset and the permuted datasets. The error bars represent the standard deviations of 550 permutations. The red dots indicate the numbers of DEGs identified from the original dataset.

**D.** Percentage of permuted datasets where a DEG identified from the original dataset was also identified as a DEG. The genes are sorted by absolute log2(fold-change) in the original dataset in decreasing order and the average values of each 100 genes are shown. The absolute log2(fold-change) values corresponding to the ranks are listed in parentheses below the ranks.The line is fitted using the loess method, and the shaded areas represent 95% confidential intervals.

**E.** GO term enrichment for the DEGs identified from at least 10% permuted datasets. The top 5 enriched biological processes GO terms are shown. The analyses were performed using R package clusterProfiler. P.adjust represents the adjusted p-value using the Benjamini & Hochberg method.

**F.** Boxplots showing the poorness of fitting the negative binomial model to the genes identified by DESeq2 or edgeR as DEGs from any permuted datasets vs. all the other genes. The poorness of fit for each gene is defined as its negative $\log_{10}$(P-value) from the Pearson's chi-squared test for the negative binomial distribution. The p-value in each panel was calculated by the Wilcoxon rank-sum test to compare the two groups of genes' poorness-of-fit values.

**Supplementary Fig. 8**

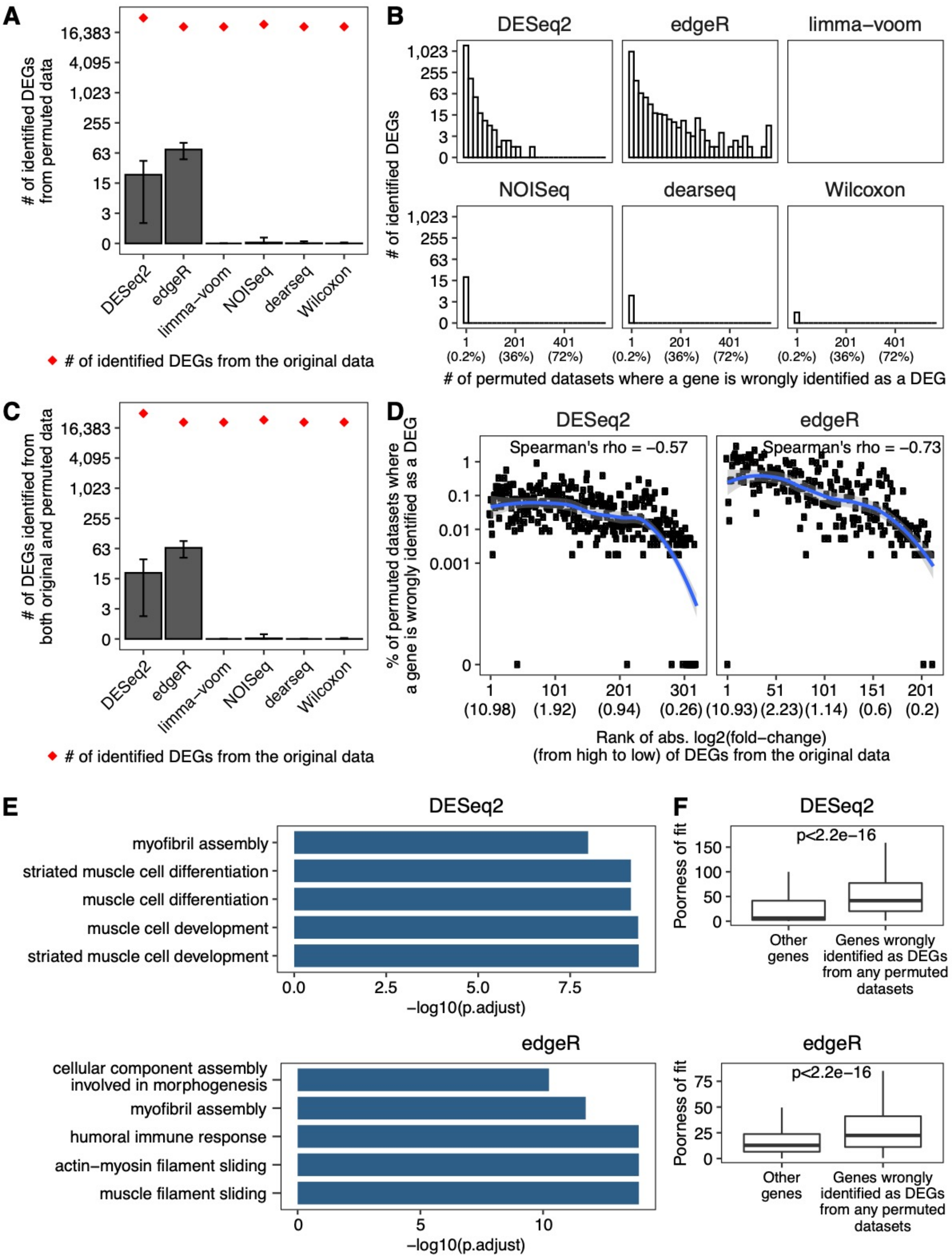


**Supplementary Fig. 8. Exaggerated false DEGs identified by DESeq2 and edgeR from prostate vs. brain cortex GTEx RNA-seq datasets.**

**A.** Barplot showing the average numbers of DEGs identified from 561 permuted datasets. The error bars represent the standard deviations of 561 permutations. The red dots indicate the numbers of DEGs identified from the original dataset.

**B.** The distributions of the number of permuted datasets where a gene was mistakenly identified as a DEG. The percentages corresponding to the numbers are listed in parentheses below the numbers.

**C.** Barplot showing the average numbers of DEGs identified from both the original dataset and the permuted datasets. The error bars represent the standard deviations of 561 permutations. The red dots indicate the numbers of DEGs identified from the original dataset.

**D.** Percentage of permuted datasets where a DEG identified from the original dataset was also identified as a DEG. The genes are sorted by absolute log2(fold-change) in the original dataset in decreasing order and the average values of each 100 genes are shown. The absolute log2(fold-change) values corresponding to the ranks are listed in parentheses below the ranks.The line is fitted using the loess method, and the shaded areas represent 95% confidential intervals.

**E.** GO term enrichment for the DEGs identified from at least 10% permuted datasets. The top 5 enriched biological processes GO terms are shown. The analyses were performed using R package clusterProfiler. P.adjust represents the adjusted p-value using the Benjamini & Hochberg method.

**F.** Boxplots showing the poorness of fitting the negative binomial model to the genes identified by DESeq2 or edgeR as DEGs from any permuted datasets vs. all the other genes. The poorness of fit for each gene is defined as its negative $\log_{10}$(P-value) from the Pearson's chi-squared test for the negative binomial distribution. The p-value in each panel was calculated by the Wilcoxon rank-sum test to compare the two groups of genes' poorness-of-fit values.

Supplementary Fig. 9

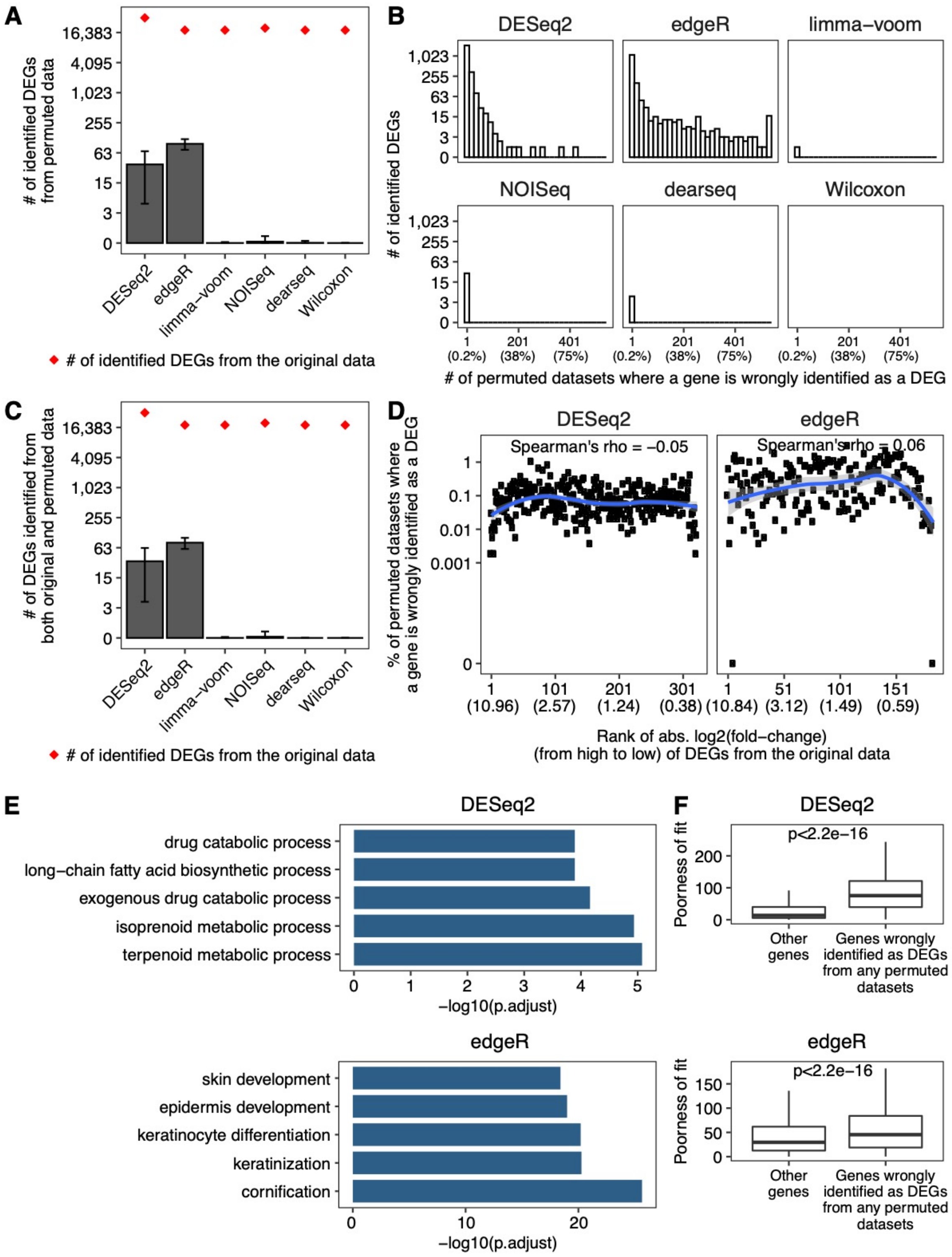


**Supplementary Fig. 9. Exaggerated false DEGs identified by DESeq2 and edgeR from whole blood vs. muscle GTEx RNA-seq datasets.**

**A.** Barplot showing the average numbers of DEGs identified from 534 permuted datasets. The error bars represent the standard deviations of 534 permutations. The red dots indicate the numbers of DEGs identified from the original dataset.

**B.** The distributions of the number of permuted datasets where a gene was mistakenly identified as a DEG. The percentages corresponding to the numbers are listed in parentheses below the numbers.

**C.** Barplot showing the average numbers of DEGs identified from both the original dataset and the permuted datasets. The error bars represent the standard deviations of 534 permutations. The red dots indicate the numbers of DEGs identified from the original dataset.

**D.** Percentage of permuted datasets where a DEG identified from the original dataset was also identified as a DEG. The genes are sorted by absolute log2(fold-change) in the original dataset in decreasing order and the average values of each 100 genes are shown. The absolute log2(fold-change) values corresponding to the ranks are listed in parentheses below the ranks.The line is fitted using the loess method, and the shaded areas represent 95% confidential intervals.

**E.** GO term enrichment for the DEGs identified from at least 10% permuted datasets. The top 5 enriched biological processes GO terms are shown. The analyses were performed using R package clusterProfiler. P.adjust represents the adjusted p-value using the Benjamini & Hochberg method.

**F.** Boxplots showing the poorness of fitting the negative binomial model to the genes identified by DESeq2 or edgeR as DEGs from any permuted datasets vs. all the other genes. The poorness of fit for each gene is defined as its negative $\log_{10}$(P-value) from the Pearson's chi-squared test for the negative binomial distribution. The p-value in each panel was calculated by the Wilcoxon rank-sum test to compare the two groups of genes' poorness-of-fit values.

Supplementary Fig. 10

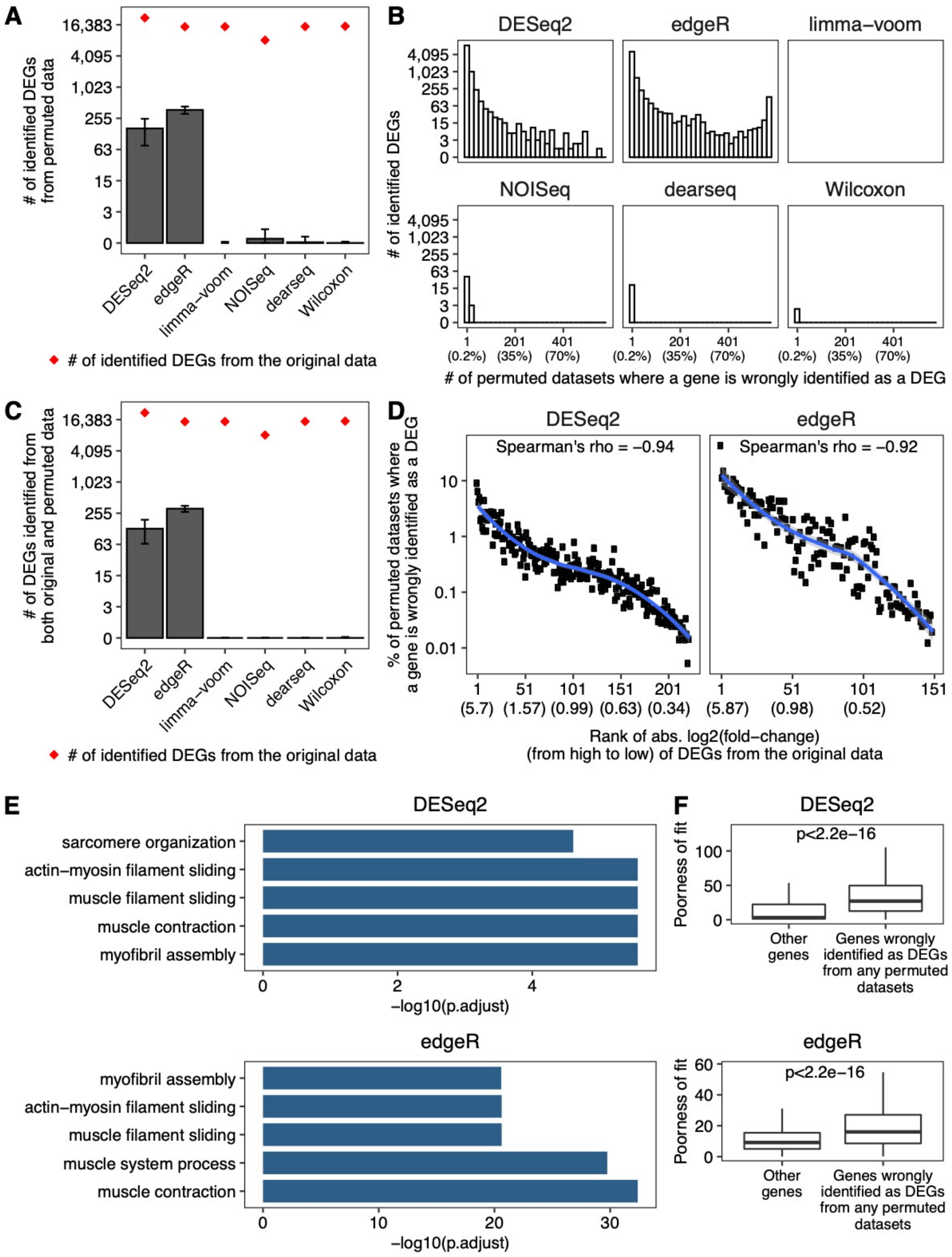


**Supplementary Fig. 10. Exaggerated false DEGs identified by DESeq2 and edgeR from BRCA (tumor vs. normal) TCGA RNA-seq datasets.**

**A.** Barplot showing the average numbers of DEGs identified from 571 permuted datasets. The error bars represent the standard deviations of 571 permutations. The red dots indicate the numbers of DEGs identified from the original dataset.

**B.** The distributions of the number of permuted datasets where a gene was mistakenly identified as a DEG. The percentages corresponding to the numbers are listed in parentheses below the numbers.

**C.** Barplot showing the average numbers of DEGs identified from both the original dataset and the permuted datasets. The error bars represent the standard deviations of 571 permutations. The red dots indicate the numbers of DEGs identified from the original dataset.

**D.** Percentage of permuted datasets where a DEG identified from the original dataset was also identified as a DEG. The genes are sorted by absolute log2(fold-change) in the original dataset in decreasing order and the average values of each 100 genes are shown. The absolute log2(fold-change) values corresponding to the ranks are listed in parentheses below the ranks.The line is fitted using the loess method, and the shaded areas represent 95% confidential intervals.

**E.** GO term enrichment for the DEGs identified from at least 10% permuted datasets. The top 5 enriched biological processes GO terms are shown. The analyses were performed using R package clusterProfiler. P.adjust represents the adjusted p-value using the Benjamini & Hochberg method.

**F.** Boxplots showing the poorness of fitting the negative binomial model to the genes identified by DESeq2 or edgeR as DEGs from any permuted datasets vs. all the other genes. The poorness of fit for each gene is defined as its negative $\log_{10}$(P-value) from the Pearson's chi-squared test for the negative binomial distribution. The p-value in each panel was calculated by the Wilcoxon rank-sum test to compare the two groups of genes' poorness-of-fit values.

**Supplementary Fig. 11**

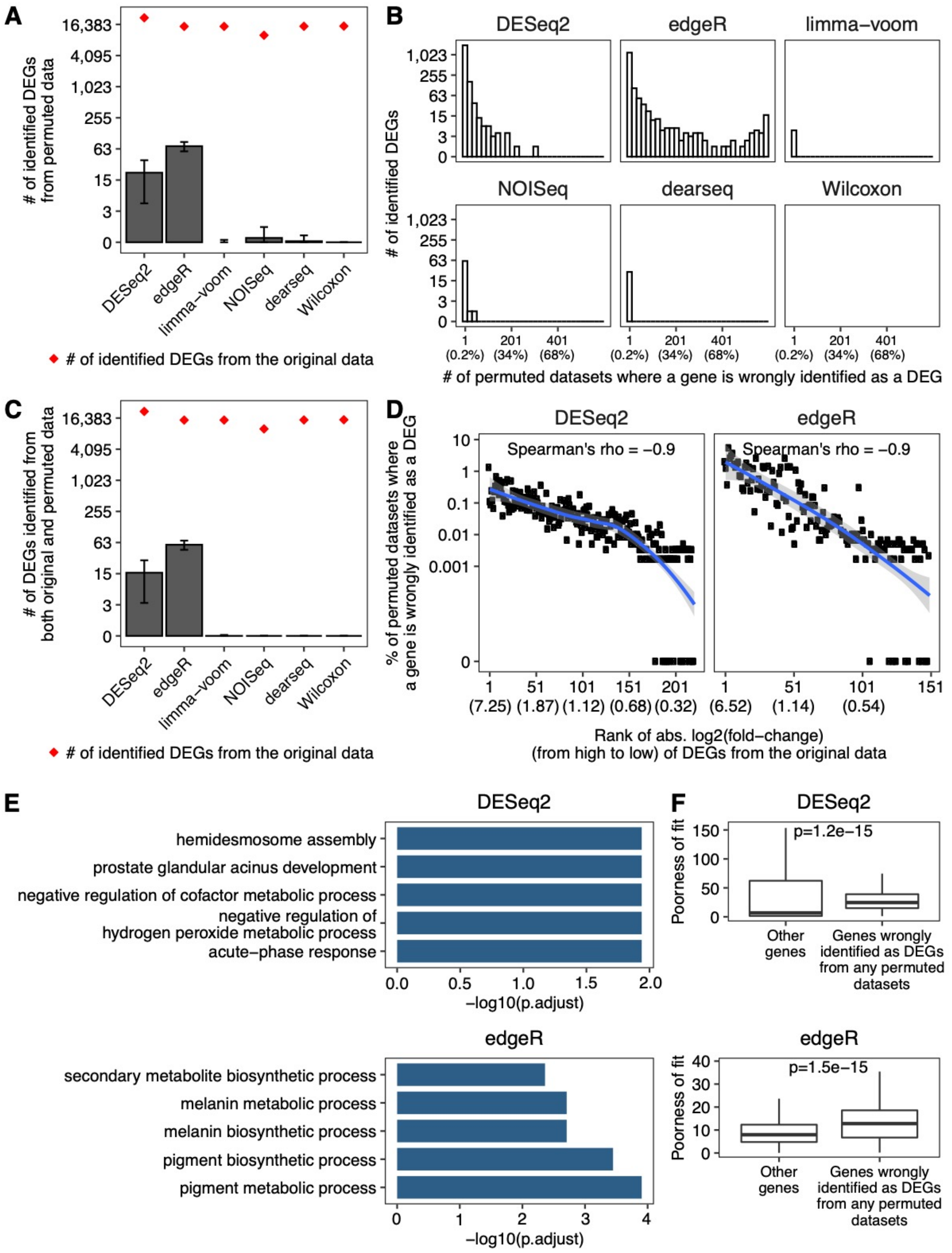


**Supplementary Fig. 11. Exaggerated false DEGs identified by DESeq2 and edgeR from KIRC (tumor vs. normal) TCGA RNA-seq datasets.**

**A.** Barplot showing the average numbers of DEGs identified from 591 permuted datasets. The error bars represent the standard deviations of 591 permutations. The red dots indicate the numbers of DEGs identified from the original dataset.

**B.** The distributions of the number of permuted datasets where a gene was mistakenly identified as a DEG. The percentages corresponding to the numbers are listed in parentheses below the numbers.

**C.** Barplot showing the average numbers of DEGs identified from both the original dataset and the permuted datasets. The error bars represent the standard deviations of 591 permutations. The red dots indicate the numbers of DEGs identified from the original dataset.

**D.** Percentage of permuted datasets where a DEG identified from the original dataset was also identified as a DEG. The genes are sorted by absolute log2(fold-change) in the original dataset in decreasing order and the average values of each 100 genes are shown. The absolute log2(fold-change) values corresponding to the ranks are listed in parentheses below the ranks.The line is fitted using the loess method, and the shaded areas represent 95% confidential intervals.

**E.** GO term enrichment for the DEGs identified from at least 10% permuted datasets. The top 5 enriched biological processes GO terms are shown. The analyses were performed using R package clusterProfiler. P.adjust represents the adjusted p-value using the Benjamini & Hochberg method.

**F.** Boxplots showing the poorness of fitting the negative binomial model to the genes identified by DESeq2 or edgeR as DEGs from any permuted datasets vs. all the other genes. The poorness of fit for each gene is defined as its negative $\log_{10}$(P-value) from the Pearson's chi-squared test for the negative binomial distribution. The p-value in each panel was calculated by the Wilcoxon rank-sum test to compare the two groups of genes' poorness-of-fit values.

**Supplementary Fig. 12**

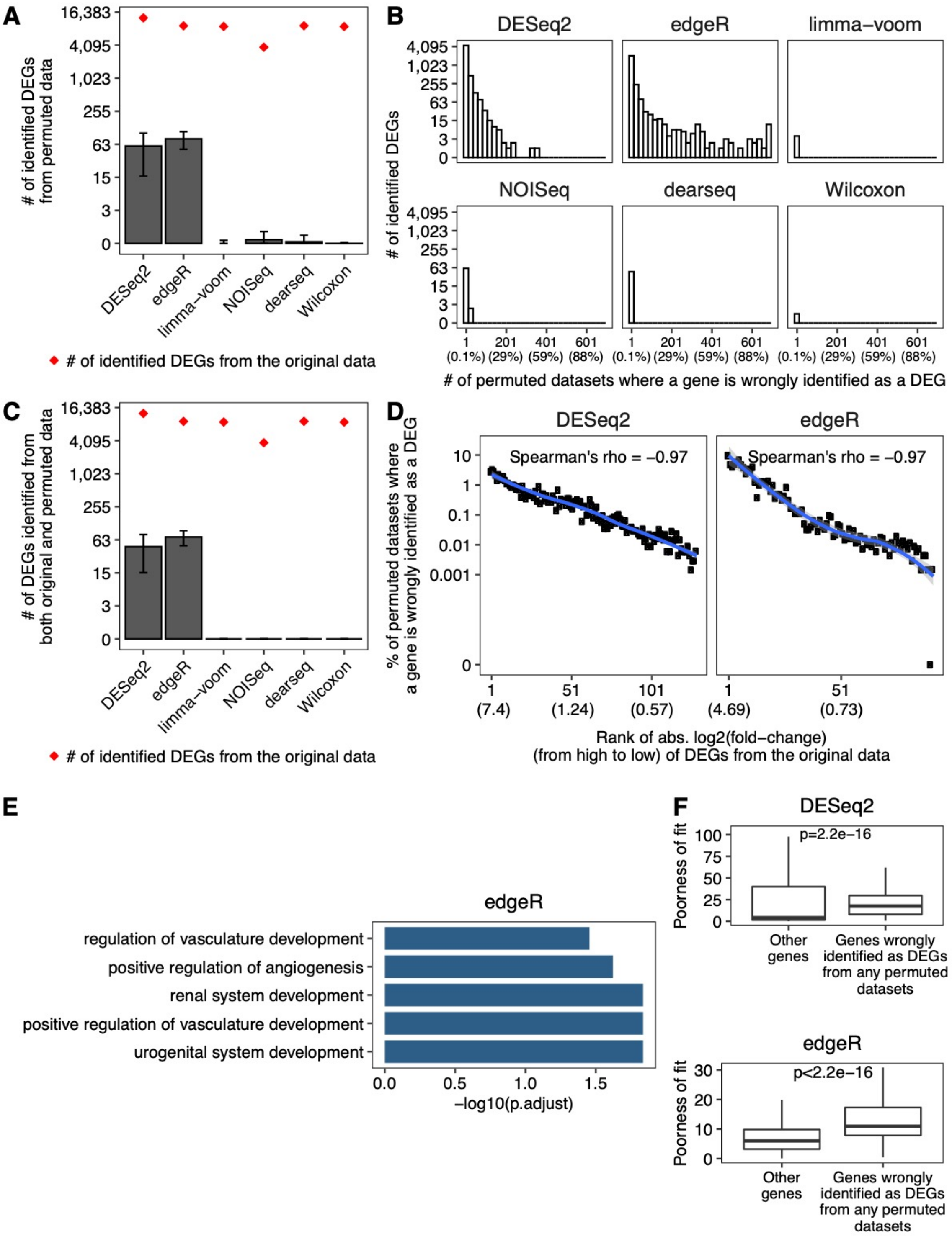


**Supplementary Fig. 12. Exaggerated false DEGs identified by DESeq2 and edgeR from LIHC (tumor vs. normal) TCGA RNA-seq datasets.**

**A.** Barplot showing the average numbers of DEGs identified from 685 permuted datasets. The error bars represent the standard deviations of 685 permutations. The red dots indicate the numbers of DEGs identified from the original dataset.

**B.** The distributions of the number of permuted datasets where a gene was mistakenly identified as a DEG. The percentages corresponding to the numbers are listed in parentheses below the numbers.

**C.** Barplot showing the average numbers of DEGs identified from both the original dataset and the permuted datasets. The error bars represent the standard deviations of 685 permutations. The red dots indicate the numbers of DEGs identified from the original dataset.

**D.** Percentage of permuted datasets where a DEG identified from the original dataset was also identified as a DEG. The genes are sorted by absolute log2(fold-change) in the original dataset in decreasing order and the average values of each 100 genes are shown. The absolute log2(fold-change) values corresponding to the ranks are listed in parentheses below the ranks.The line is fitted using the loess method, and the shaded areas represent 95% confidential intervals.

**E.** GO term enrichment for the DEGs identified from at least 10% permuted datasets. The top 5 enriched biological processes GO terms are shown. The analyses were performed using R package clusterProfiler. P.adjust represents the adjusted p-value using the Benjamini & Hochberg method.

**F.** Boxplots showing the poorness of fitting the negative binomial model to the genes identified by DESeq2 or edgeR as DEGs from any permuted datasets vs. all the other genes. The poorness of fit for each gene is defined as its negative $\log_{10}$(P-value) from the Pearson's chi-squared test for the negative binomial distribution. The p-value in each panel was calculated by the Wilcoxon rank-sum test to compare the two groups of genes' poorness-of-fit values.

**Supplementary Fig. 13**

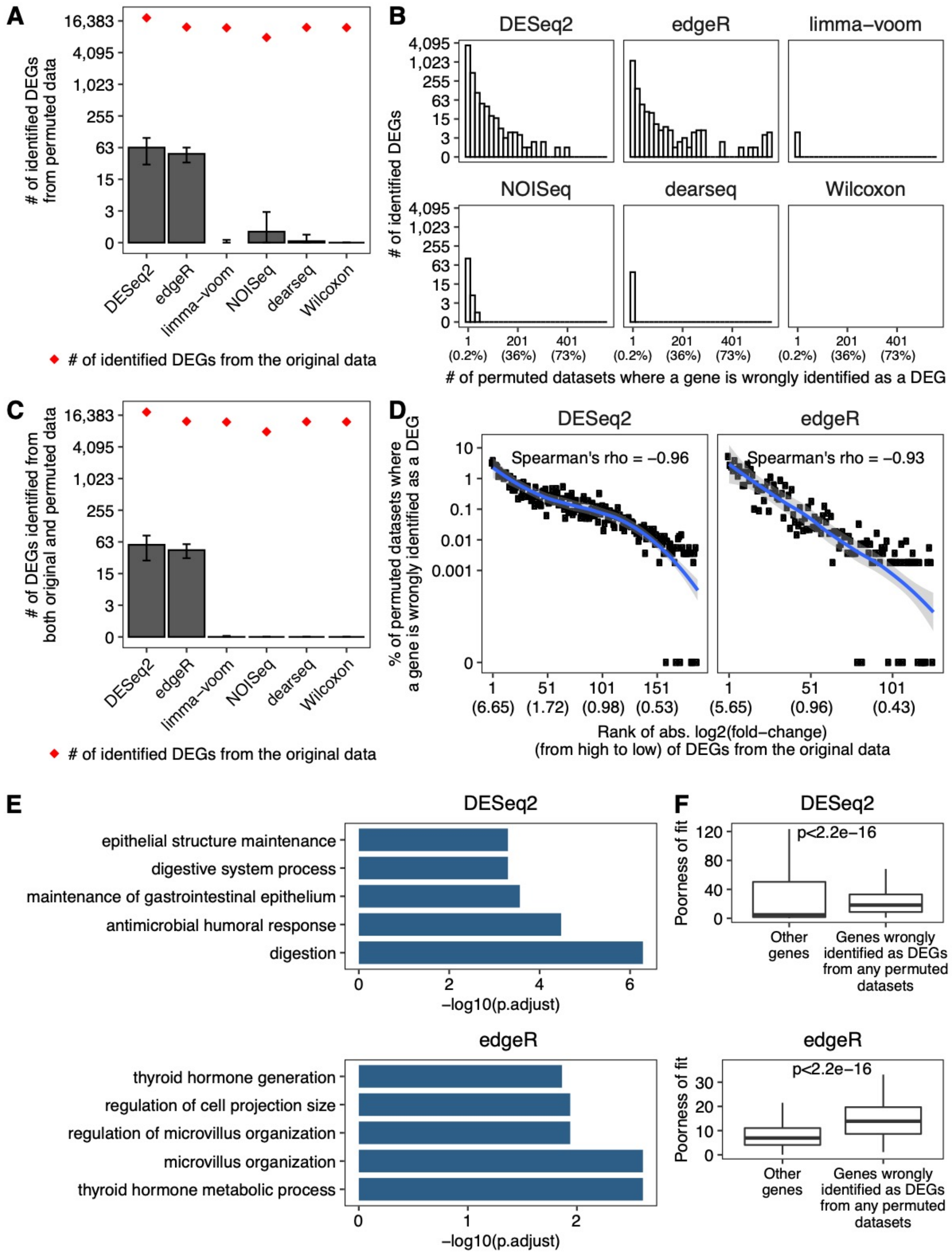


**Supplementary Fig. 13. Exaggerated false DEGs identified by DESeq2 and edgeR from LUAD (tumor vs. normal) TCGA RNA-seq datasets.**

**A.** Barplot showing the average numbers of DEGs identified from 552 permuted datasets. The error bars represent the standard deviations of 552 permutations. The red dots indicate the numbers of DEGs identified from the original dataset.

**B.** The distributions of the number of permuted datasets where a gene was mistakenly identified as a DEG. The percentages corresponding to the numbers are listed in parentheses below the numbers.

**C.** Barplot showing the average numbers of DEGs identified from both the original dataset and the permuted datasets. The error bars represent the standard deviations of 552 permutations. The red dots indicate the numbers of DEGs identified from the original dataset.

**D.** Percentage of permuted datasets where a DEG identified from the original dataset was also identified as a DEG. The genes are sorted by absolute log2(fold-change) in the original dataset in decreasing order and the average values of each 100 genes are shown. The absolute log2(fold-change) values corresponding to the ranks are listed in parentheses below the ranks.The line is fitted using the loess method, and the shaded areas represent 95% confidential intervals.

**E.** GO term enrichment for the DEGs identified from at least 10% permuted datasets. The top 5 enriched biological processes GO terms are shown. The analyses were performed using R package clusterProfiler. P.adjust represents the adjusted p-value using the Benjamini & Hochberg method.

**F.** Boxplots showing the poorness of fitting the negative binomial model to the genes identified by DESeq2 or edgeR as DEGs from any permuted datasets vs. all the other genes. The poorness of fit for each gene is defined as its negative $\log_{10}$(P-value) from the Pearson's chi-squared test for the negative binomial distribution. The p-value in each panel was calculated by the Wilcoxon rank-sum test to compare the two groups of genes' poorness-of-fit values.

**Supplementary Fig. 14**

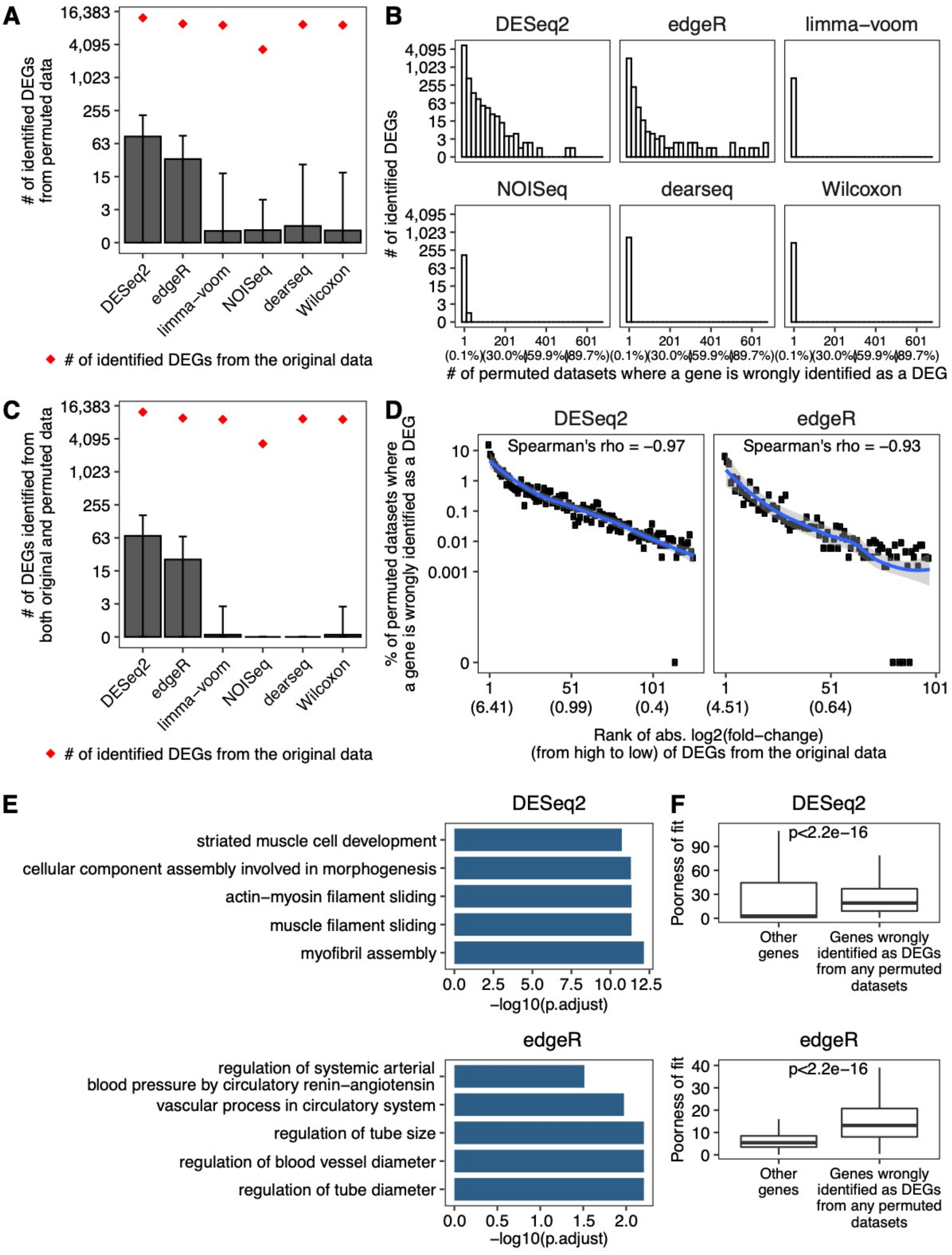


**Supplementary Fig. 14. Exaggerated false DEGs identified by DESeq2 and edgeR from PRAD (tumor vs. normal) TCGA RNA-seq datasets.**

**A.** Barplot showing the average numbers of DEGs identified from 670 permuted datasets. The error bars represent the standard deviations of 670 permutations. The red dots indicate the numbers of DEGs identified from the original dataset.

**B.** The distributions of the number of permuted datasets where a gene was mistakenly identified as a DEG. The percentages corresponding to the numbers are listed in parentheses below the numbers.

**C.** Barplot showing the average numbers of DEGs identified from both the original dataset and the permuted datasets. The error bars represent the standard deviations of 670 permutations. The red dots indicate the numbers of DEGs identified from the original dataset.

**D.** Percentage of permuted datasets where a DEG identified from the original dataset was also identified as a DEG. The genes are sorted by absolute log2(fold-change) in the original dataset in decreasing order and the average values of each 100 genes are shown. The absolute log2(fold-change) values corresponding to the ranks are listed in parentheses below the ranks.The line is fitted using the loess method, and the shaded areas represent 95% confidential intervals.

**E.** GO term enrichment for the DEGs identified from at least 10% permuted datasets. The top 5 enriched biological processes GO terms are shown. The analyses were performed using R package clusterProfiler. P.adjust represents the adjusted p-value using the Benjamini & Hochberg method.

**F.** Boxplots showing the poorness of fitting the negative binomial model to the genes identified by DESeq2 or edgeR as DEGs from any permuted datasets vs. all the other genes. The poorness of fit for each gene is defined as its negative $\log_{10}$(P-value) from the Pearson's chi-squared test for the negative binomial distribution. The p-value in each panel was calculated by the Wilcoxon rank-sum test to compare the two groups of genes' poorness-of-fit values.

**Supplementary Fig. 15**

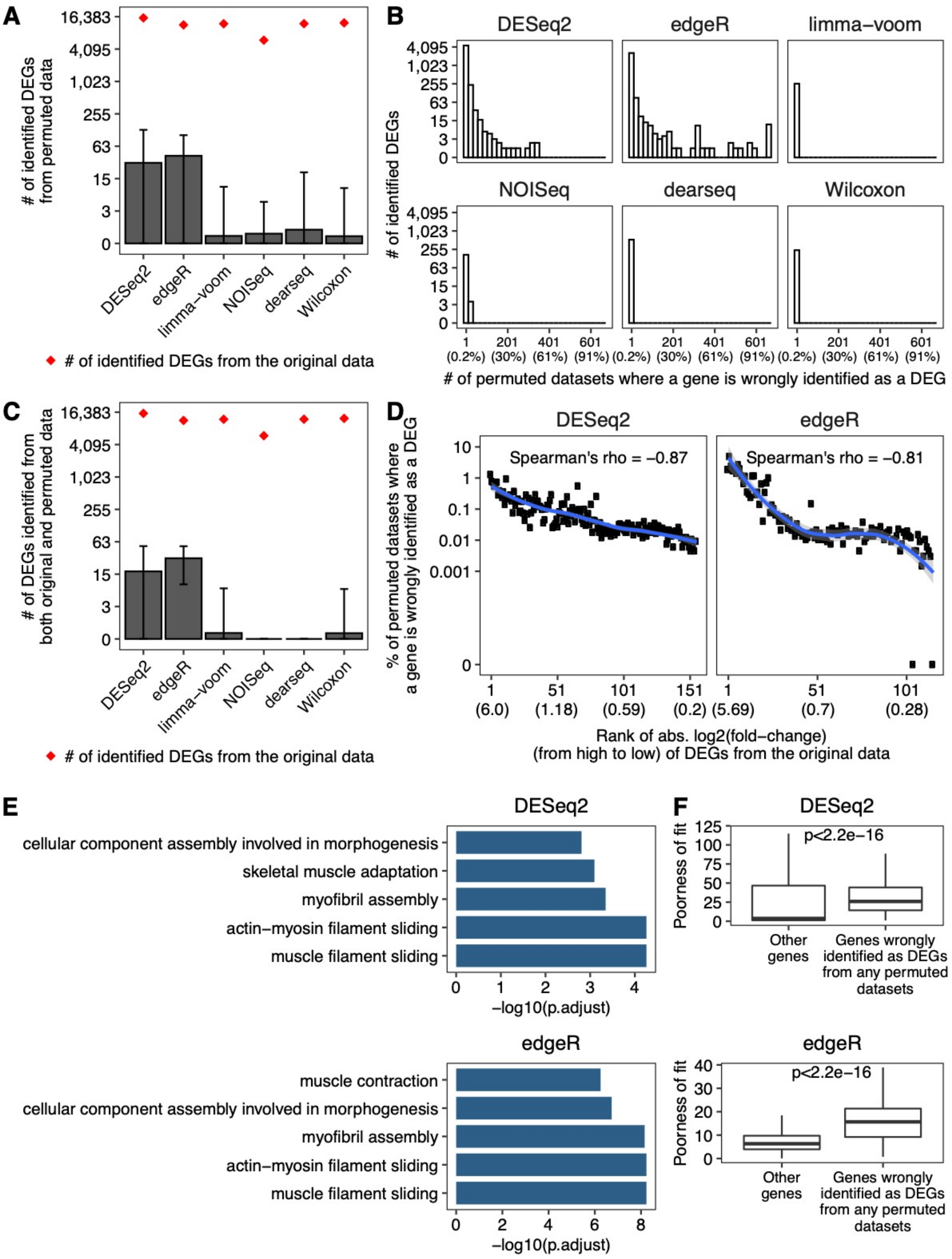


**Supplementary Fig. 15. Exaggerated false DEGs identified by DESeq2 and edgeR from THCA (tumor vs. normal) TCGA RNA-seq datasets.**

**A.** Barplot showing the average numbers of DEGs identified from 662 permuted datasets. The error bars represent the standard deviations of 662 permutations. The red dots indicate the numbers of DEGs identified from the original dataset.

**B.** The distributions of the number of permuted datasets where a gene was mistakenly identified as a DEG. The percentages corresponding to the numbers are listed in parentheses below the numbers.

**C.** Barplot showing the average numbers of DEGs identified from both the original dataset and the permuted datasets. The error bars represent the standard deviations of 662 permutations. The red dots indicate the numbers of DEGs identified from the original dataset.

**D.** Percentage of permuted datasets where a DEG identified from the original dataset was also identified as a DEG. The genes are sorted by absolute log2(fold-change) in the original dataset in decreasing order and the average values of each 100 genes are shown. The absolute log2(fold-change) values corresponding to the ranks are listed in parentheses below the ranks.The line is fitted using the loess method, and the shaded areas represent 95% confidential intervals.

**E.** GO term enrichment for the DEGs identified from at least 10% permuted datasets. The top 5 enriched biological processes GO terms are shown. The analyses were performed using R package clusterProfiler. P.adjust represents the adjusted p-value using the Benjamini & Hochberg method.

**F.** Boxplots showing the poorness of fitting the negative binomial model to the genes identified by DESeq2 or edgeR as DEGs from any permuted datasets vs. all the other genes. The poorness of fit for each gene is defined as its negative $\log_{10}$(P-value) from the Pearson's chi-squared test for the negative binomial distribution. The p-value in each panel was calculated by the Wilcoxon rank-sum test to compare the two groups of genes' poorness-of-fit values.

**Supplementary Fig. 16**

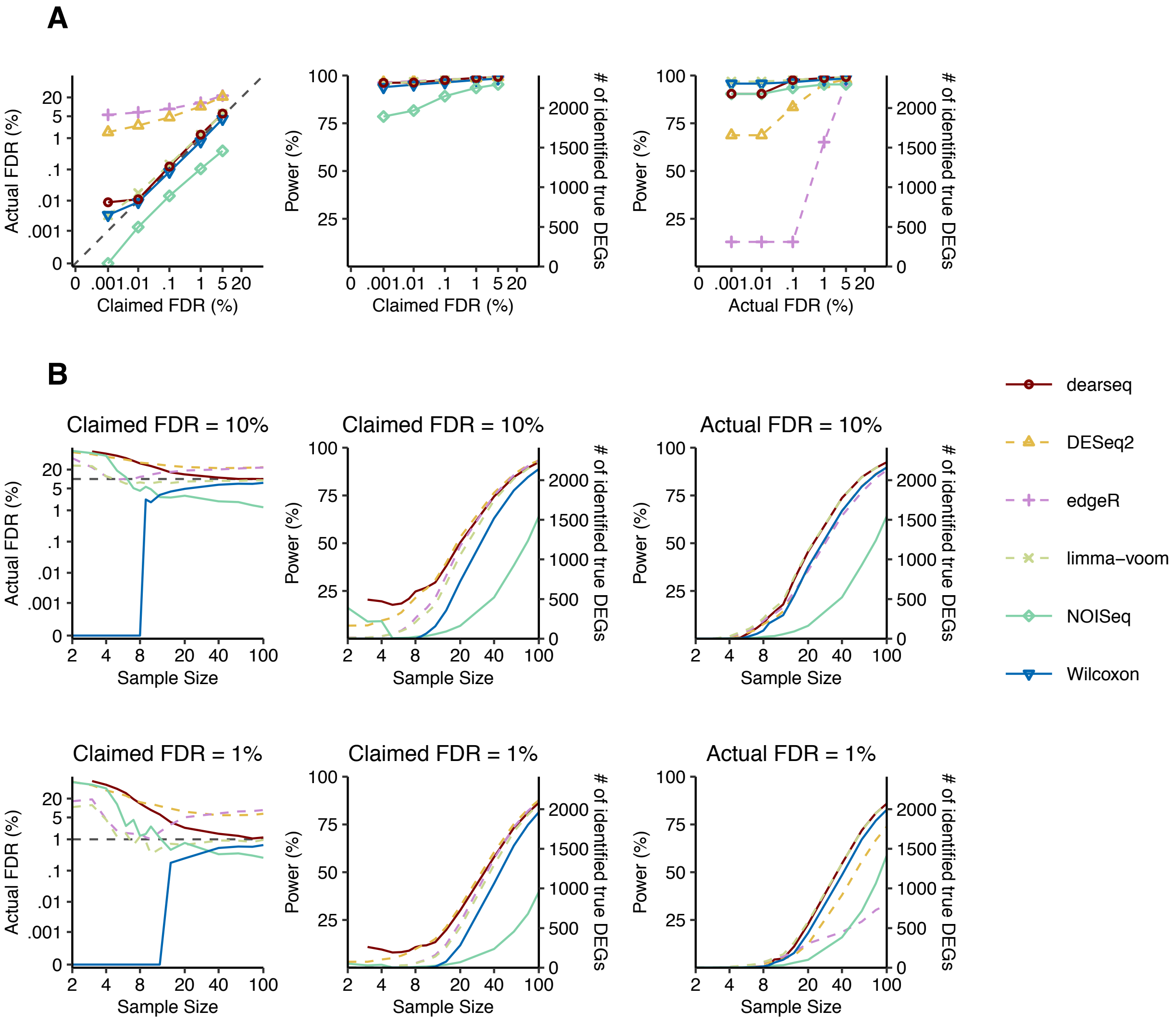


**Supplementary Fig. 16. Wilcoxon test has the best FDR control and power on adipose (subcutaneous vs. visceral) GTEx datasets with synthetic ground truths.**

**A.** The FDR control (left panel), power (middle panel) given the claimed FDRs, and power given the actual FDRs (right panel) under a range of FDR thresholds from 0.001% to 5%.

**B.** The FDR control (left), power given the claimed FDRs (middle), and power given the actual FDRs (right) for a range of per-condition sample sizes from 2 to 100, under FDR thresholds 10% (top panels) and 1% (bottom panels). The claimed FDRs, actual FDRs, and power were all calculated as the averages of 50 randomly down-sampled datasets.

### Supplementary Fig. 17

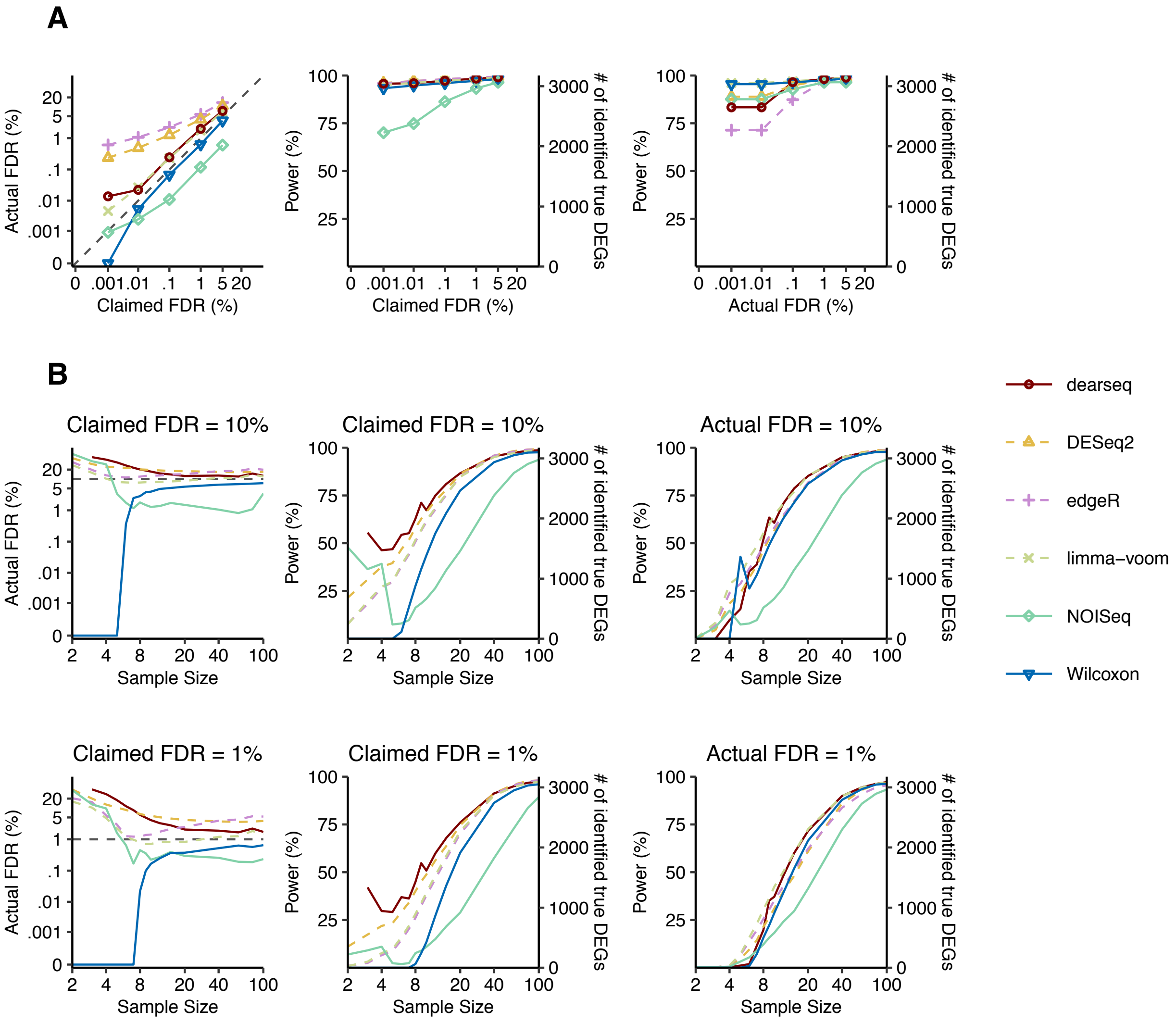


**Supplementary Fig. 17. Wilcoxon test has the best FDR control and power on brain (amygdala vs. spinal cord) GTEx datasets with synthetic ground truths.**

**A.** The FDR control (left panel), power (middle panel) given the claimed FDRs, and power given the actual FDRs (right panel) under a range of FDR thresholds from 0.001% to 5%.

**B.** The FDR control (left), power given the claimed FDRs (middle), and power given the actual FDRs (right) for a range of per-condition sample sizes from 2 to 100, under FDR thresholds 10% (top panels) and 1% (bottom panels). The claimed FDRs, actual FDRs, and power were all calculated as the averages of 50 randomly down-sampled datasets.

### Supplementary Fig. 18

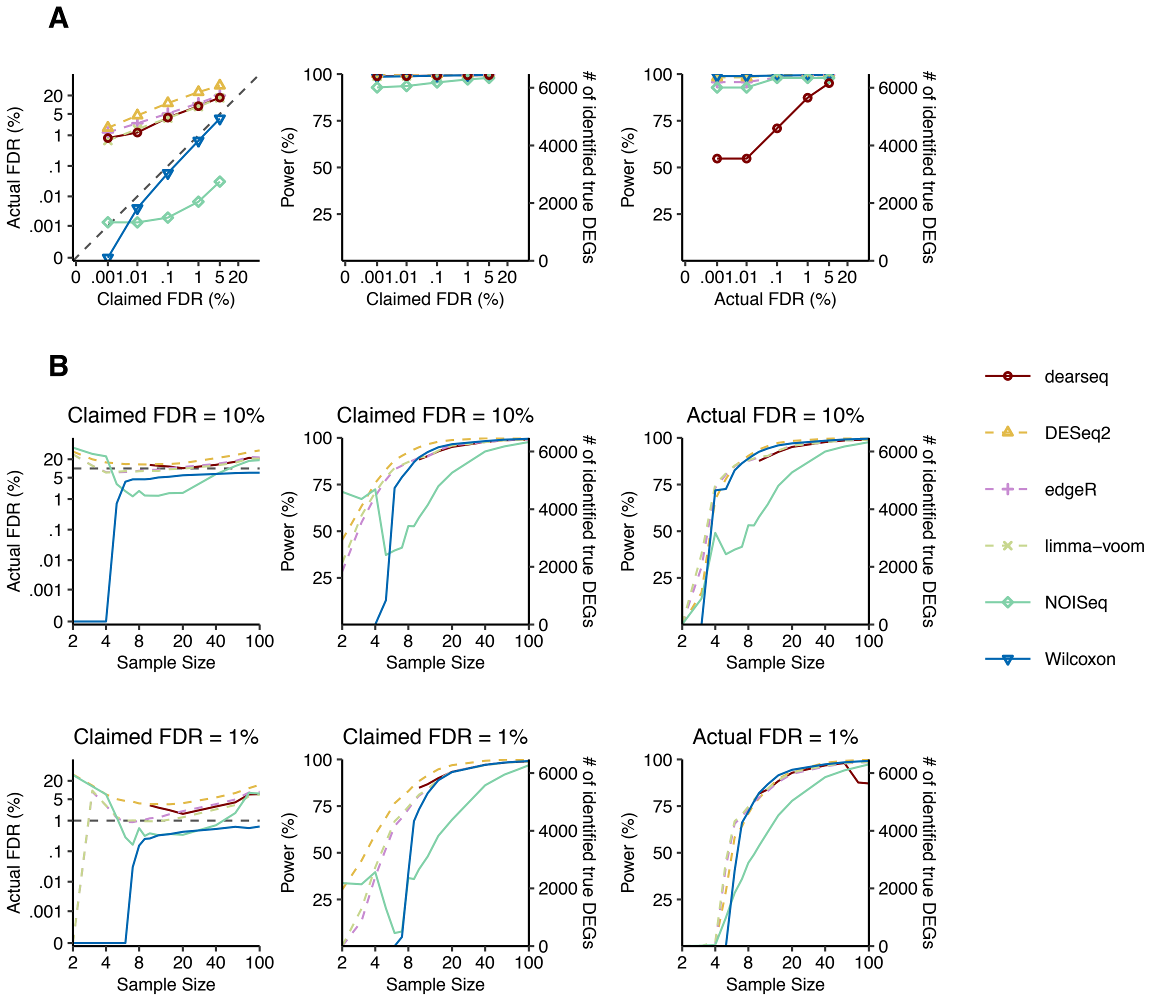


**Supplementary Fig. 18. Wilcoxon test has the best FDR control and power on EVB transformed lymphocytes vs. minor salivary gland GTEx datasets with synthetic ground truths.**

**A.** The FDR control (left panel), power (middle panel) given the claimed FDRs, and power given the actual FDRs (right panel) under a range of FDR thresholds from 0.001% to 5%.

**B.** The FDR control (left), power given the claimed FDRs (middle), and power given the actual FDRs (right) for a range of per-condition sample sizes from 2 to 100, under FDR thresholds 10% (top panels) and 1% (bottom panels). The claimed FDRs, actual FDRs, and power were all calculated as the averages of 50 randomly down-sampled datasets.

### Supplementary Fig. 19

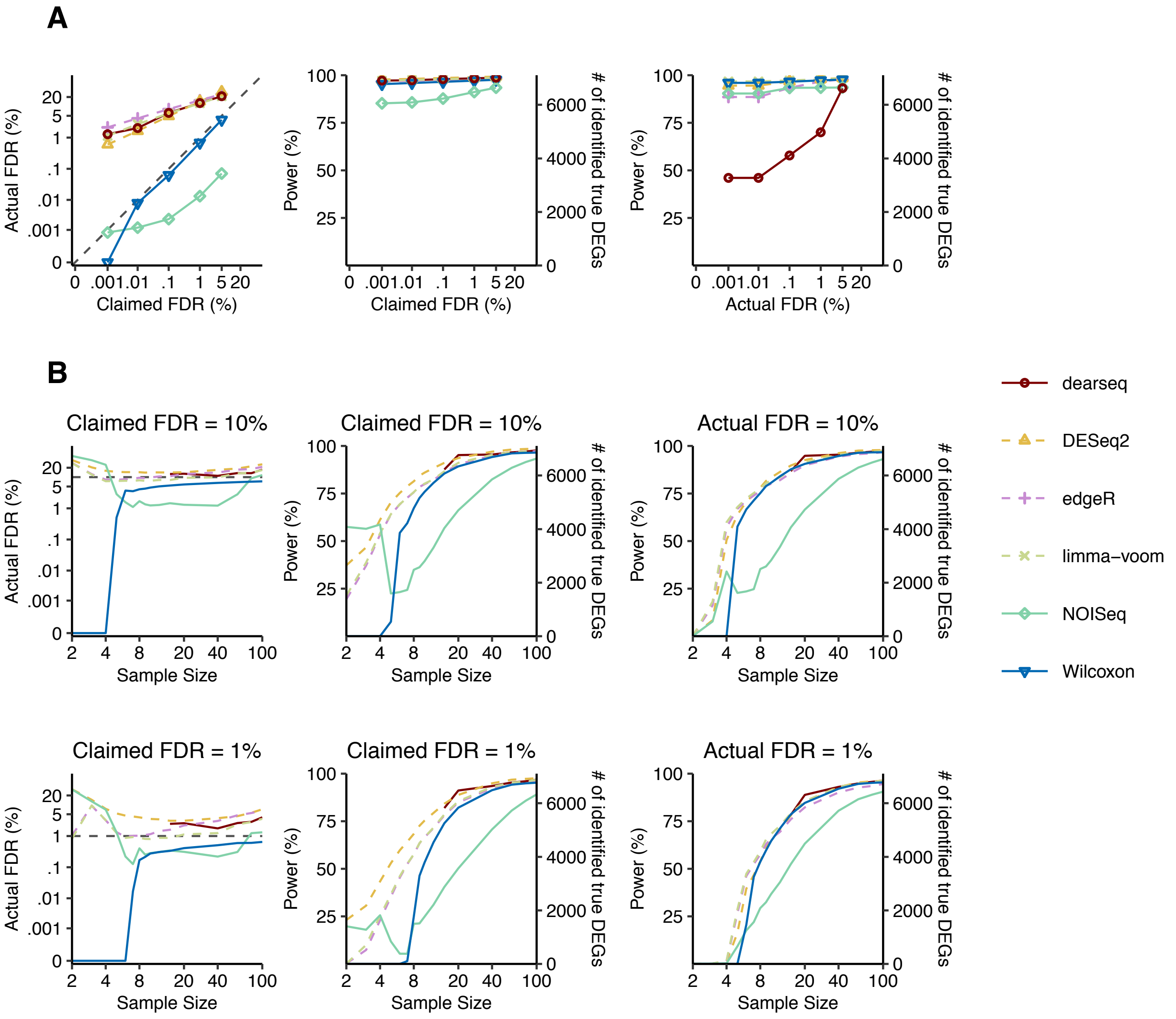


**Supplementary Fig. 19. Wilcoxon test has the best FDR control and power on prostate vs. brain cortex GTEx datasets with synthetic ground truths.**

**A.** The FDR control (left panel), power (middle panel) given the claimed FDRs, and power given the actual FDRs (right panel) under a range of FDR thresholds from 0.001% to 5%.

**B.** The FDR control (left), power given the claimed FDRs (middle), and power given the actual FDRs (right) for a range of per-condition sample sizes from 2 to 100, under FDR thresholds 10% (top panels) and 1% (bottom panels). The claimed FDRs, actual FDRs, and power were all calculated as the averages of 50 randomly down-sampled datasets.

**Supplementary Fig. 20**

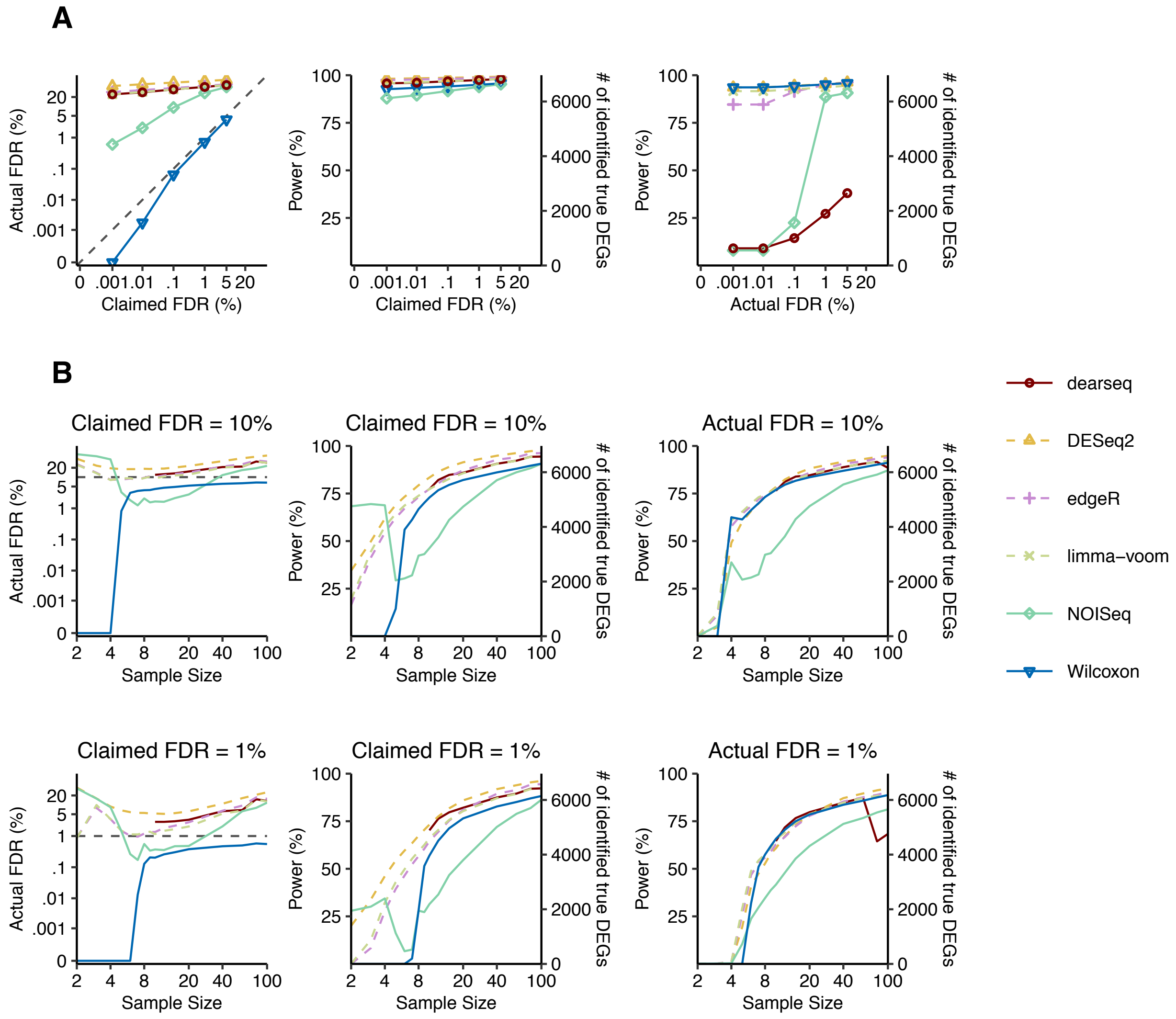


**Supplementary Fig. 20. Wilcoxon test has the best FDR control and power on whole blood vs. muscle GTEx datasets with synthetic ground truths.**

**A.** The FDR control (left panel), power (middle panel) given the claimed FDRs, and power given the actual FDRs (right panel) under a range of FDR thresholds from 0.001% to 5%.

**B.** The FDR control (left), power given the claimed FDRs (middle), and power given the actual FDRs (right) for a range of per-condition sample sizes from 2 to 100, under FDR thresholds 10% (top panels) and 1% (bottom panels). The claimed FDRs, actual FDRs, and power were all calculated as the averages of 50 randomly down-sampled datasets.

### Supplementary Fig. 21

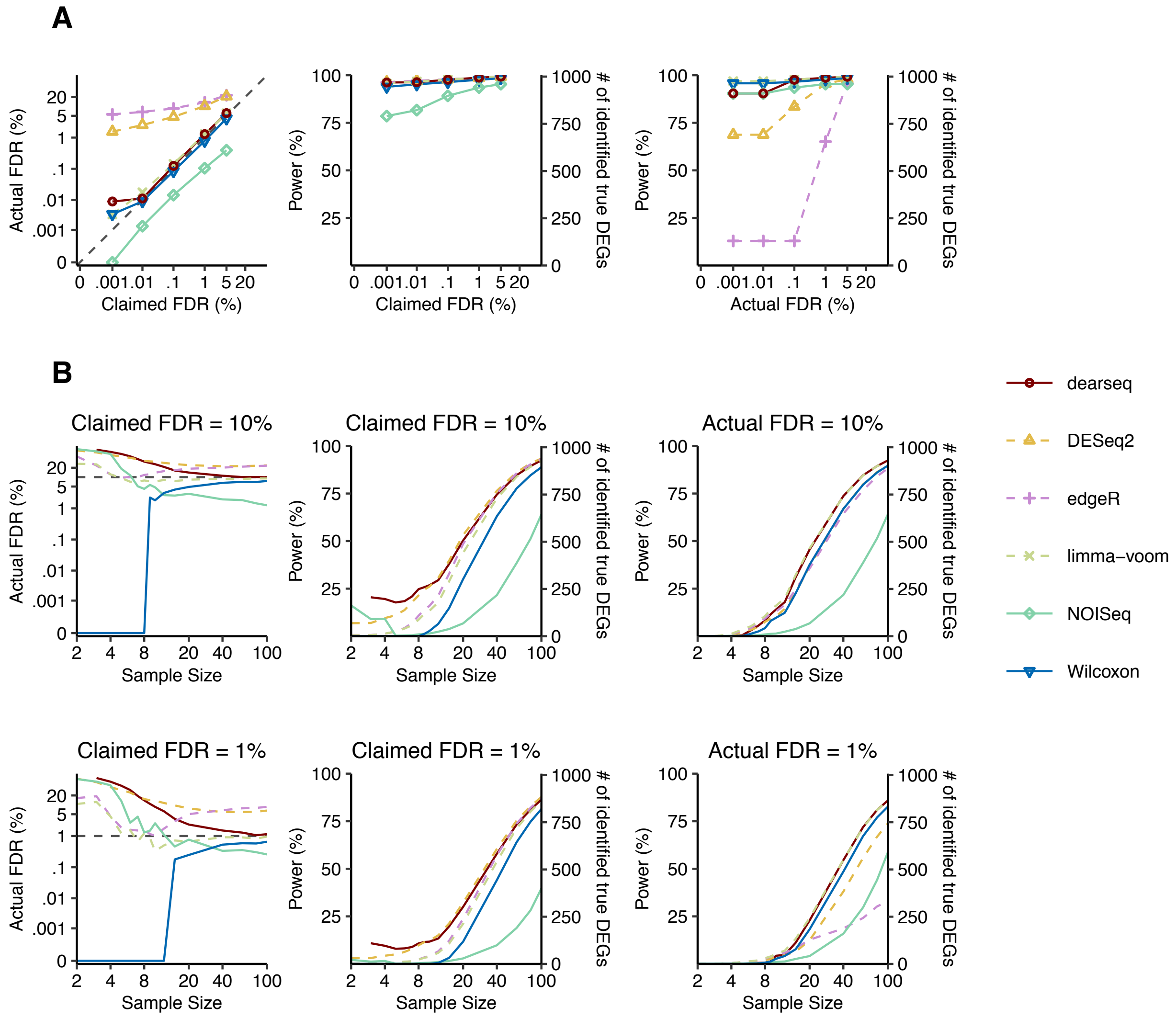


**Supplementary Fig. 21. Wilcoxon test has the best FDR control and power on BRCA TCGA datasets with synthetic ground truths.**

**A.** The FDR control (left panel), power (middle panel) given the claimed FDRs, and power given the actual FDRs (right panel) under a range of FDR thresholds from 0.001% to 5%.

**B.** The FDR control (left), power given the claimed FDRs (middle), and power given the actual FDRs (right) for a range of per-condition sample sizes from 2 to 100, under FDR thresholds 10% (top panels) and 1% (bottom panels). The claimed FDRs, actual FDRs, and power were all calculated as the averages of 50 randomly down-sampled datasets.

**Supplementary Fig. 22**

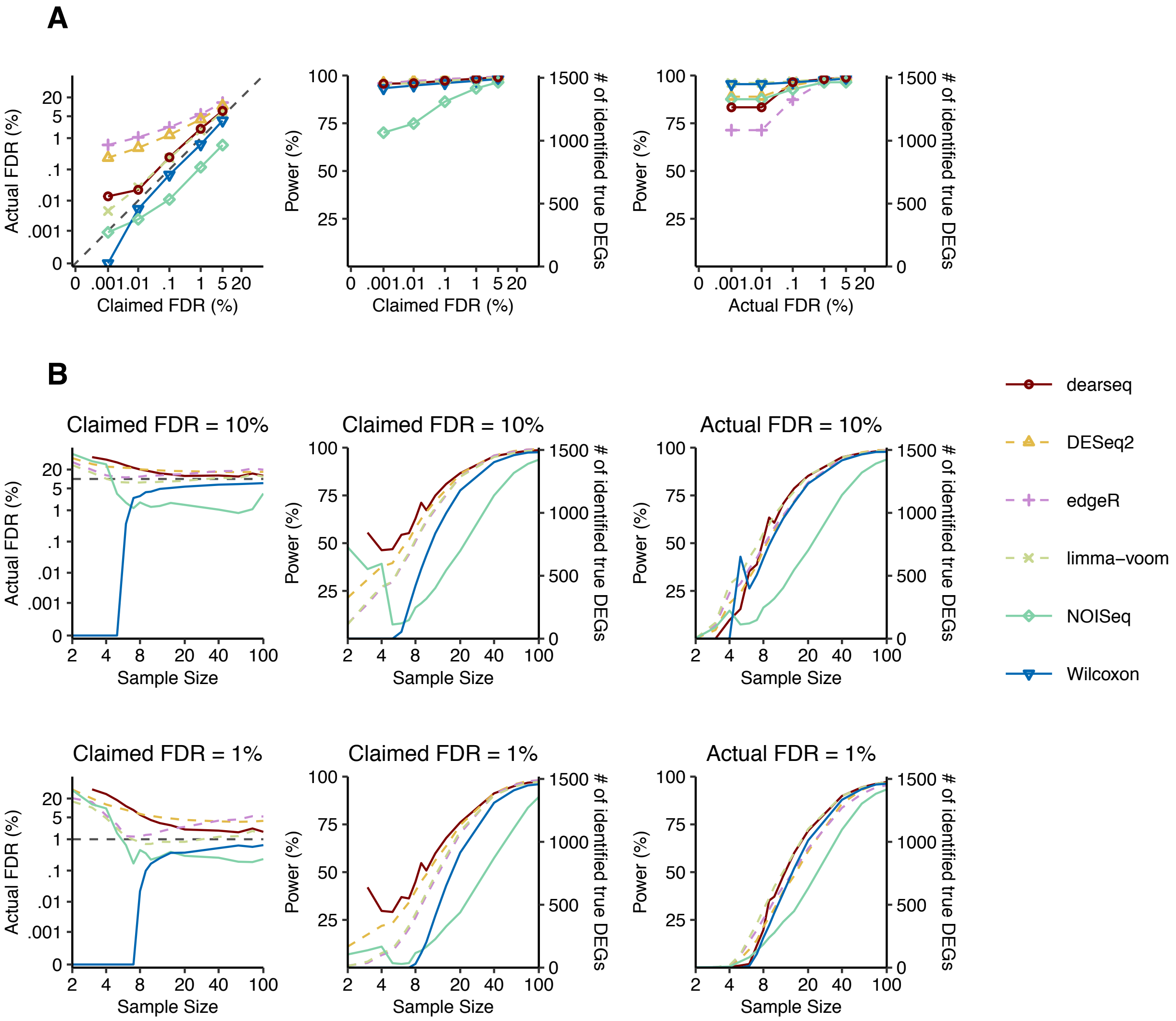


**Supplementary Fig. 22. Wilcoxon test has the best FDR control and power on KIRC TCGA datasets with synthetic ground truths.**

**A.** The FDR control (left panel), power (middle panel) given the claimed FDRs, and power given the actual FDRs (right panel) under a range of FDR thresholds from 0.001% to 5%.

**B.** The FDR control (left), power given the claimed FDRs (middle), and power given the actual FDRs (right) for a range of per-condition sample sizes from 2 to 100, under FDR thresholds 10% (top panels) and 1% (bottom panels). The claimed FDRs, actual FDRs, and power were all calculated as the averages of 50 randomly down-sampled datasets.

**Supplementary Fig. 23**

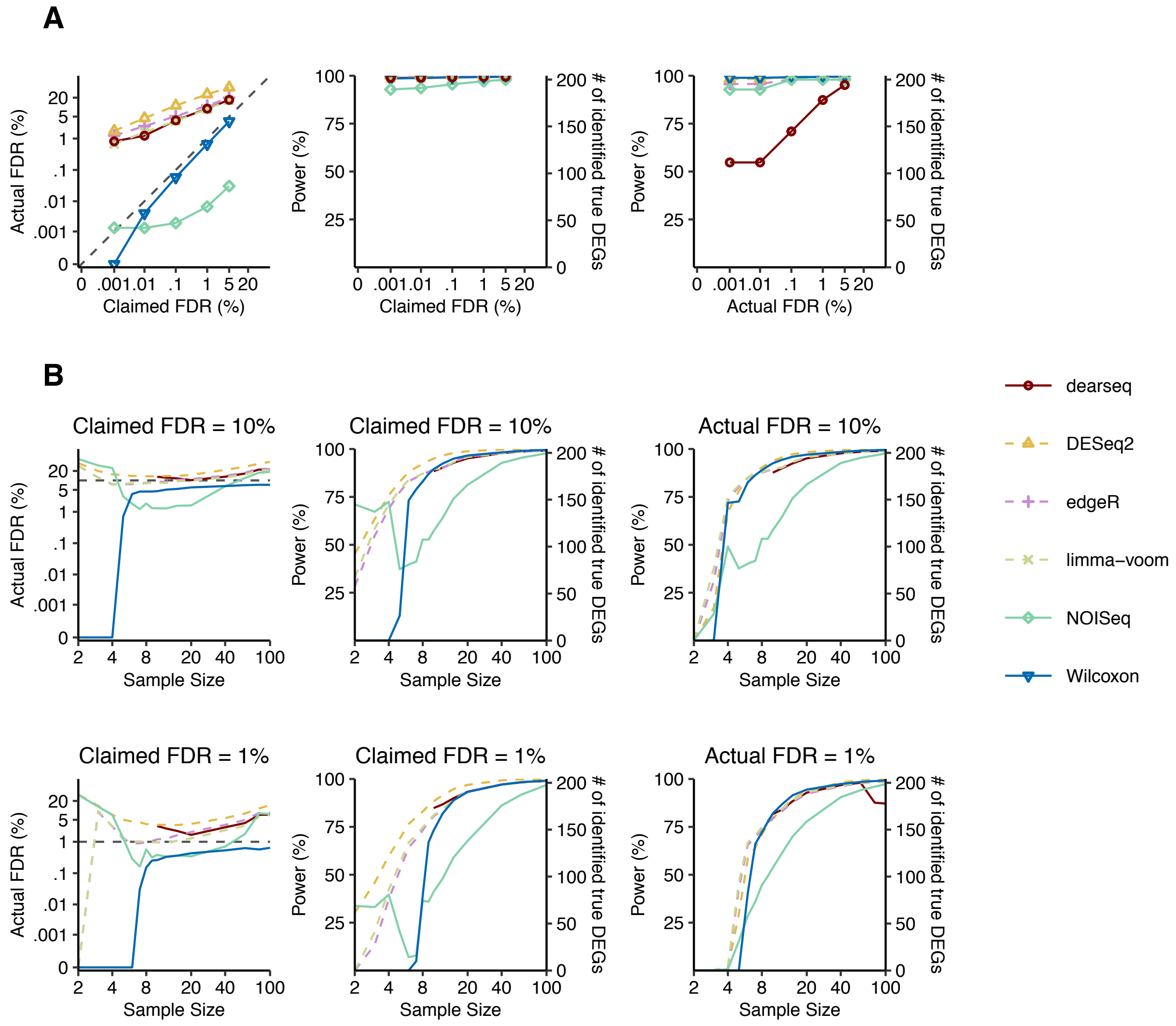


**Supplementary Fig. 23. Wilcoxon test has the best FDR control and power on LIHC TCGA datasets with synthetic ground truths.**

**A.** The FDR control (left panel), power (middle panel) given the claimed FDRs, and power given the actual FDRs (right panel) under a range of FDR thresholds from 0.001% to 5%.

**B.** The FDR control (left), power given the claimed FDRs (middle), and power given the actual FDRs (right) for a range of per-condition sample sizes from 2 to 100, under FDR thresholds 10% (top panels) and 1% (bottom panels). The claimed FDRs, actual FDRs, and power were all calculated as the averages of 50 randomly down-sampled datasets.

**Supplementary Fig. 24**

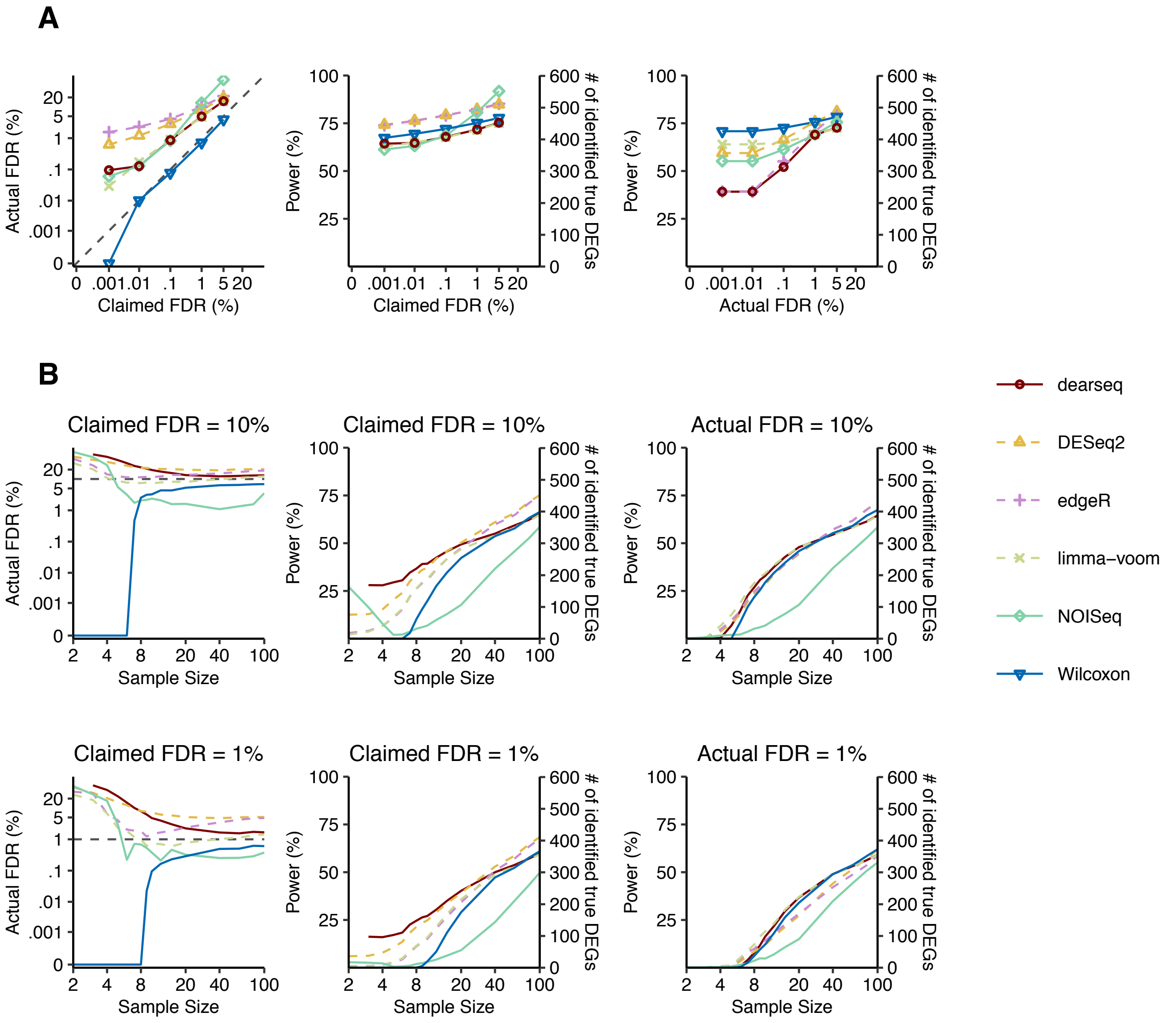


**Supplementary Fig. 24. Wilcoxon test has the best FDR control and power on LUAD TCGA datasets with synthetic ground truths.**

**A.** The FDR control (left panel), power (middle panel) given the claimed FDRs, and power given the actual FDRs (right panel) under a range of FDR thresholds from 0.001% to 5%.

**B.** The FDR control (left), power given the claimed FDRs (middle), and power given the actual FDRs (right) for a range of per-condition sample sizes from 2 to 100, under FDR thresholds 10% (top panels) and 1% (bottom panels). The claimed FDRs, actual FDRs, and power were all calculated as the averages of 50 randomly down-sampled datasets.

### Supplementary Fig. 25

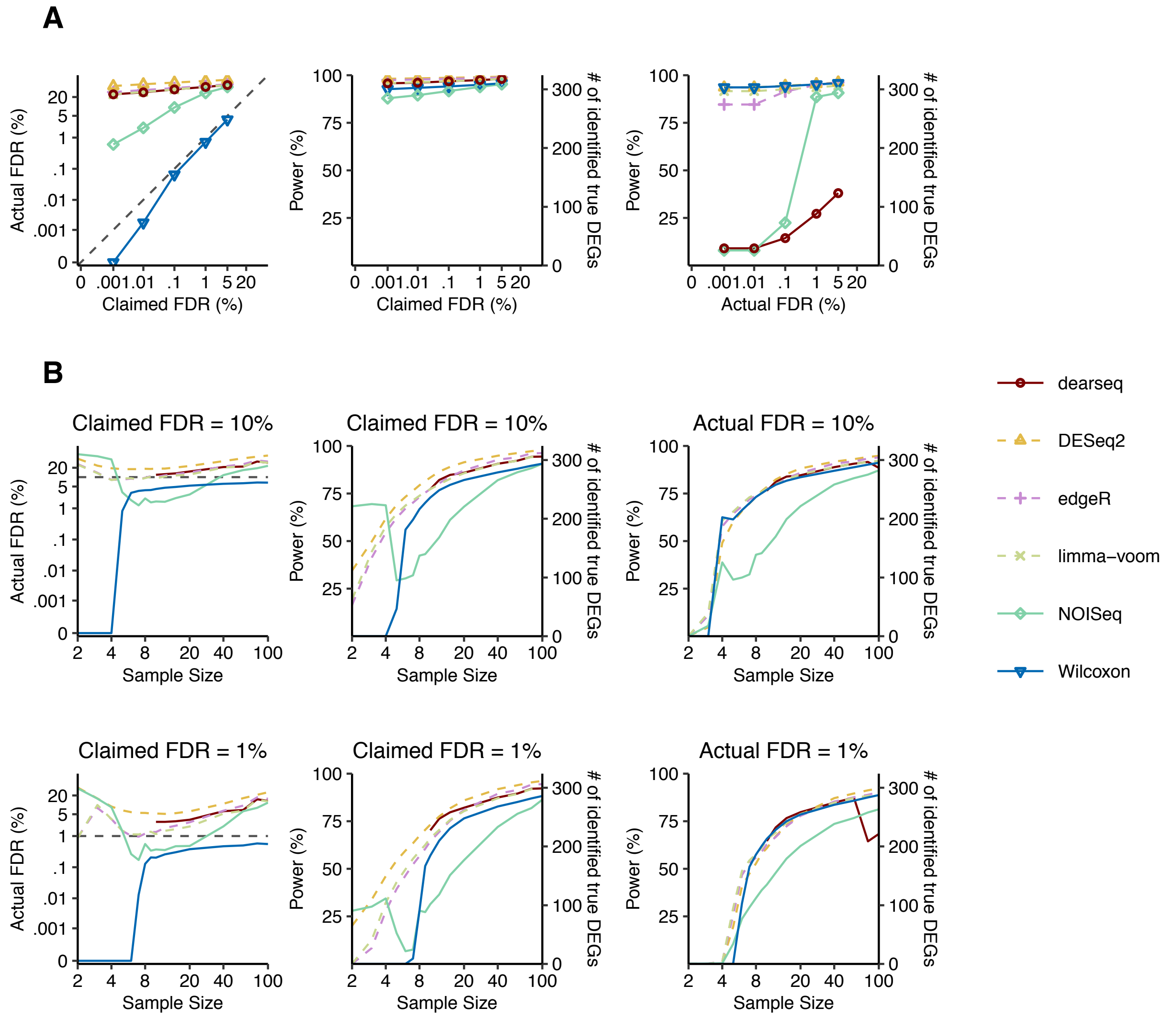


**Supplementary Fig. 25. Wilcoxon test has the best FDR control and power on PRAD TCGA datasets with synthetic ground truths.**

**A.** The FDR control (left panel), power (middle panel) given the claimed FDRs, and power given the actual FDRs (right panel) under a range of FDR thresholds from 0.001% to 5%.

**B.** The FDR control (left), power given the claimed FDRs (middle), and power given the actual FDRs (right) for a range of per-condition sample sizes from 2 to 100, under FDR thresholds 10% (top panels) and 1% (bottom panels). The claimed FDRs, actual FDRs, and power were all calculated as the averages of 50 randomly down-sampled datasets.

**Supplementary Fig. 26**

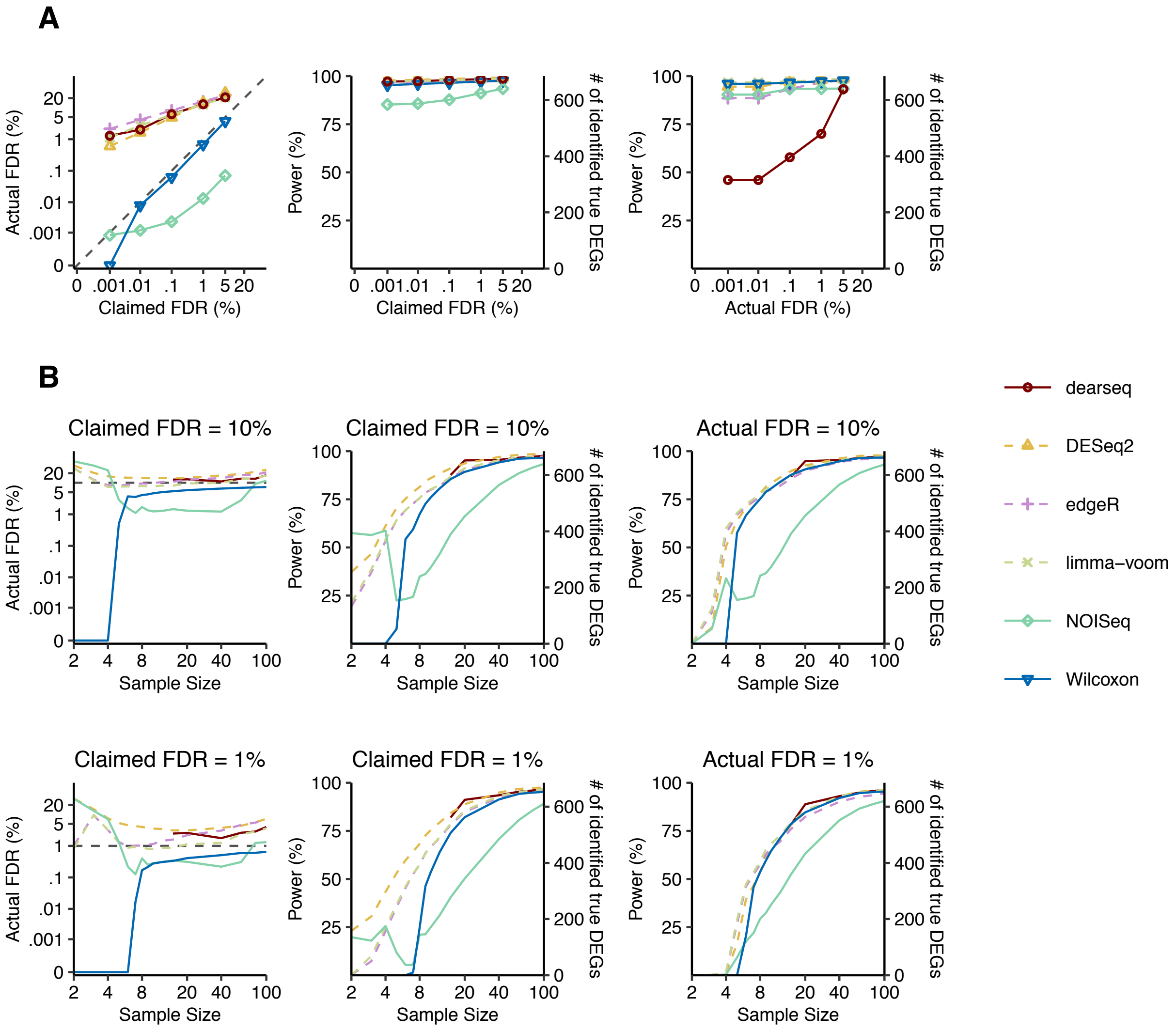


**Supplementary Fig. 26. Wilcoxon test has the best FDR control and power on THCA TCGA datasets with synthetic ground truths.**

**A.** The FDR control (left panel), power (middle panel) given the claimed FDRs, and power given the actual FDRs (right panel) under a range of FDR thresholds from 0.001% to 5%.

**B.** The FDR control (left), power given the claimed FDRs (middle), and power given the actual FDRs (right) for a range of per-condition sample sizes from 2 to 100, under FDR thresholds 10% (top panels) and 1% (bottom panels). The claimed FDRs, actual FDRs, and power were all calculated as the averages of 50 randomly down-sampled datasets.

### Supplementary Table 1. Summary of information for samples used in this study

| Data source | Condition 1 | Condition 2 | Sample size (Condition 1 vs. Condition 2) | Accession |
|---|---|---|---|---|
| **Immunotherapy study** | Pre-therapy | On-therapy | 51 vs. 58 | GSE91061 |
| **TCGA** | BRCA normal tissue[1] | BRCA tumor tissue | 112 vs. 112 | GDC Xena Hub |
| | KIRC normal tissue[2] | KIRC tumor tissue | 72 vs. 72 | |
| | THCA normal tissue[3] | THCA tumor tissue | 58 vs. 58 | |
| | LUAD normal tissue[4] | LUAD tumor tissue | 57 vs. 57 | |
| | PRAD normal tissue[5] | PRAD tumor tissue | 52 vs. 52 | |
| | LIHC normal tissue[6] | LIHC tumor tissue | 50 vs. 50 | |
| **GTEx** | Whole blood | Muscle - skeletal | 670 vs. 706 | GTEx Portal |
| | Adipose - subcutaneous | Adipose - visceral | 581 vs. 469 | |
| | Heart - atrial appendage | Heart - Left ventricle | 372 vs. 386 | |
| | Prostate | Brain - cortex | 221 vs. 205 | |
| | Cells - EVB transformed lymphocytes | Minor salivary gland | 147 vs. 144 | |
| | Brain - amygdala | Brain - spinal cord | 129 vs. 126 | |

[1]BRCA: Breast invasive carcinoma

[2]KIRC: Kidney renal clear cell carcinoma

[3]THCA: Thyroid carcinoma

[4]LUAD: Lung adenocarcinoma

[5]PRAD: Prostate adenocarcinoma

[6]LIHC: Liver hepatocellular carcinoma
